# SRSF1 shapes 3′-end site selection with differential dependence on U1 snRNP

**DOI:** 10.64898/2026.04.01.715904

**Authors:** Hope E. Merens, Ana-Maria Raicu, Christine L. Carroll, Mike Kourkoulakos, Ana Fiszbein, L. Stirling Churchman

## Abstract

Proper polyadenylation site (PAS) selection is critical for RNA isoform determination. Core spliceosomal components, including U1 snRNP, regulate PAS choice, but whether they work with other splicing factors in this role remains unclear. Here, we establish that the splicing factor SRSF1 regulates 3′-end selection through complementary mechanisms with varying degrees of independence from U1 snRNP. Within 3’ UTRs, largely independent of U1 snRNP, SRSF1 binds RNA near proximal PASs and promotes their usage. Congruent with this observation, breast cancer tumors with altered SRSF1 levels display shifted 3′-end selection. At PASs modulated by both SRSF1 and U1 snRNP, SRSF1 acts on sites through U1 snRNP-mediated Pol II interactions. Consistent with co-transcriptional regulation, SRSF1 reduces the Pol II elongation index and transcription readthrough. Together, our results reveal that SRSF1 shapes RNA isoform determination beyond its canonical role in splicing, through a combination of direct RNA binding and U1 snRNP-dependent co-transcriptional coordination.

## Introduction

To become mature and translatable, eukaryotic pre-mRNA must be transcribed and properly processed. Two major steps of pre-mRNA processing are splicing and cleavage and polyadenylation (CPA). The core splicing machinery – including the essential U1, U2, U4, U5, and U6 snRNPs – is necessary for constitutive splicing. To generate alternative splicing patterns, additional splicing factors are deployed, including the serine-arginine (SR) proteins, SRSF1 through SRSF12. In a manner analogous to alternative splicing, alternative polyadenylation site (PAS) selection is a dynamic choice between competing 3’-end sites^1,2^, and is shaped by both the core CPA machinery and additional regulatory factors. Together, alternative splicing and polyadenylation determine the fate of transcripts, jointly influencing RNA stability, localization, and translation, expanding the diversity of protein isoforms produced^1,3–7^. Furthermore, misregulated splicing and PAS selection are both observed in, and thought to contribute to, a number of diseases, including neurodegenerative diseases, premature aging diseases, and cancer^5,8,9^.

A substantial amount of splicing and CPA occurs co-transcriptionally^10–14^. Pre-mRNA processing factors, including splicing machinery, are enriched in RNA Polymerase II (Pol II) interactomes, as observed in yeast and human cells^15–17^. These interactions are hypothesized to promote efficient coupling between transcription and processing; in support of this model, a number of factors that interact with Pol II, such as SR proteins, stimulate co-transcriptional splicing^15,18,19^. Importantly, co-transcriptional processing is also influenced by transcription dynamics. Pol II elongation kinetically competes with splicing^20–24^, CPA, and transcription termination^24–28^, influencing the probabilities of which alternative RNA processing events will be selected. Therefore, factors that associate with Pol II and affect its elongation rate have the potential to influence PAS choice not only by directly interacting with the RNA processing machinery, but also by modulating transcription dynamics.

Among the best characterized examples of factors with co-transcriptional roles beyond their canonical functions is U1 snRNP, the component of the early spliceosome complex responsible for 5’ splice site recognition. U1 snRNP co-transcriptionally inhibits widespread intronic premature cleavage and polyadenylation (PCPA) ^29–31^, and cryo-EM structures of U1 snRNP bound to nascent RNA and to Pol II^32^ suggest that direct interactions between U1 snRNP and Pol II could allow U1 snRNP to inhibit intronic PAS usage through the regulation of transcription dynamics. Indeed, U1 snRNP accelerates Pol II elongation^33^ and suppresses transcription termination^34^. Beyond U1 snRNP, core spliceosomal snRNPs and RNA binding proteins also influence PAS selection through diverse mechanisms^35–39^. Whether U1 snRNP acts alone or in concert with other factors to shape Pol II dynamics and PAS choice remains an open question.

In this study, we asked whether SRSF1, a SR protein that uniquely interacts with both Pol II and U1 snRNP^15,40–42^, coordinates with U1 snRNP to couple transcription to PAS selection. Using a combination of genomic and biochemical approaches in human cell lines, we demonstrate that SRSF1 shapes PAS usage at hundreds of sites. By comparing the effects of SRSF1 and U1 snRNP depletion on PAS selection, we identify sites that are predominantly sensitive to SRSF1 and sites where both factors converge, revealing complementary mechanisms with varying degrees of U1 snRNP dependence. At PASs predominantly sensitive to SRSF1, which are enriched in 3’ UTRs, SRSF1 binds RNA towards the start of the 3’ UTR to promote nearby PAS usage. Moreover, the relationship we observe between SRSF1 and PAS usage is recapitulated in breast cancer tumors with altered SRSF1 levels. At sites where SRSF1 and U1 snRNP converge, SRSF1 acts on PAS usage through U1 snRNP-dependent SRSF1–Pol II interactions, and we show more broadly that U1 snRNP mediates interactions between select factors and the transcription machinery. Consistent with a kinetic model of co-transcriptional PAS selection through Pol II interactions, SRSF1 reduces the average Pol II elongation index and limits transcription readthrough, linking its effects on PAS selection to the regulation of transcription dynamics. Together, these results reveal that SRSF1 shapes 3’-end processing through complementary mechanisms — direct RNA binding and U1 snRNP-dependent coordination with Pol II — placing it at the intersection of splicing, transcription, and 3’-end processing.

## Results

### SRSF1 regulates PAS selection

To study the effect of SRSF1 on PAS usage, we used an shRNA to deplete SRSF1 for 72 hours in HEK293T cells, and collected chromatin-associated RNA (**Fig. S1A-S1B**). By analyzing chromatin-associated RNA, we sought to identify changes in PAS usage in newly transcribed transcripts before post-transcriptional turnover could obscure them. Next, we performed 3′-end sequencing that identifies the location of polyadenylated 3′-ends in RNA (**Fig. 1A, S1C-S1D**). Depletion of SRSF1 led to genome-wide changes in PAS selection, affecting the relative usage of 1284 PASs (p-adjusted < 0.05, log_2_(fold change) > |log_2_(1.5)|) out of 13,092 total identified PASs in our dataset (**Fig. 1B**). Specifically, we identified 514 significantly upregulated events and 770 significantly downregulated events. As an example, knockdown (KD) of SRSF1 led to increased usage of a proximal PAS (i.e. upstream PAS, closer to the transcription start site (TSS)) relative to a distal PAS (i.e. downstream PAS, farther from the TSS) in the 3′ UTR of *ANAPC7* (**Fig. 1B**-**1C**). In contrast, in the 3′ UTR of *IFNAR1,* SRSF1 KD resulted in decreased usage of a proximal PAS (**Fig. 1D**). Using 3′ RACE, we validated the shifts in PAS selection identified through 3′-end sequencing for the genes *ANAPC7* and *IFNAR1*, recapitulating these results in whole cell RNA (**Fig. 1E-1F**). The majority (∼68%) of significantly affected sites were in annotated 3′ UTRs (**Fig. 1G**, **S1E**). Nevertheless, across all genomic features, SRSF1 KD led to increased distal PAS usage (**Fig. 1H**). Together, these results establish SRSF1 as a widespread regulator of PAS selection that promotes the usage of proximal PASs across diverse genomic features.

**Figure 1.**
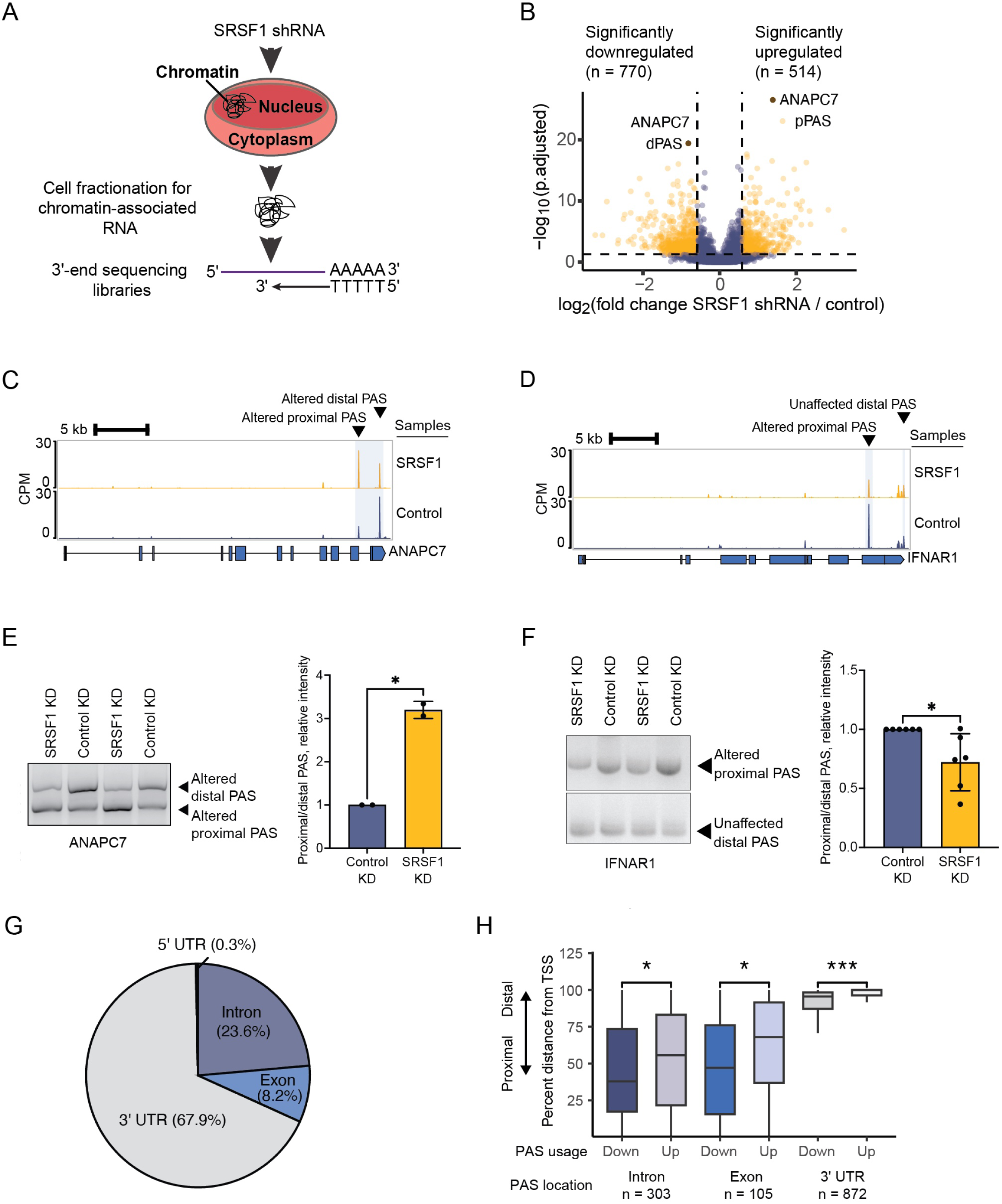
SRSF1 regulates PAS selection. A) Schematic for the chromatin-associated RNA 3′-end sequencing experiment. B) Volcano plot of the log_2_ fold change in relative PAS usage upon SRSF1 KD, as calculated by DEXseq (n = 3 replicates). Each dot is a PAS. Yellow dots are significantly changed PASs, purple dots are unaffected PASs, and brown dots represent a particular example shown later in the figure. 514 sites had significantly upregulated usage, 770 sites had significantly downregulated usage, and 11,808 sites were unchanged (p.adjusted < 0.05, abs(log_2_(fold change)) > log_2_(1.5). The distal and proximal PASs (dPAS and pPAS respectively) highlighted in C for *ANAPC7* are marked. C-D) Genome browser tracks of chromatin-associated RNA 3′-end sequencing data for genes *ANAPC7* (C) and *IFNAR1* (D) upon SRSF1 or control KD. E-F) 3′ RACE assays performed on whole cell RNA upon SRSF1 or control KD (left) and quantification (right) for *ANAPC7* (E, n = 2) and *IFNAR1* (F, n = 6; 2 replicates shown in gel). G) Significantly changed PASs shown in B mapped to their genomic features. H) Boxplots of the percent distance from the TSS to the most distal PAS detected in our 3′-end sequencing experiment for significantly altered PASs upon SRSF1 KD. The percent distance for each PAS was examined based on the direction of the altered usage (down vs. upregulated) and the genomic feature the PAS mapped to. Outlier datapoints not shown.

#### SRSF1 and U1 snRNP regulate a shared set of PASs

As SRSF1 interacts with U1 snRNP in the context of splicing^15,41–43^, we asked whether they act together in PAS selection. We performed U1-70K shRNA KD in HEK293T cells for 72 hours, and collected chromatin-associated RNA for 3′-end sequencing to compare to our SRSF1 KD data (**Fig. S2A-S2C**). Our 3′-end sequencing results identified 1924 significant changes in PAS usage upon U1-70K KD out of 16,541 total identified PASs. Specifically, there were 1,060 significantly upregulated PASs, and 864 significantly downregulated PASs (**Fig. 2A**). We compared affected sites from the U1-70K and SRSF1 KDs and found that, out of the 12,283 common PASs profiled in both datasets, many of these sites were uniquely significantly altered in one perturbation or the other (**Fig. 2B**). 937 sites were specific to SRSF1 KD (U1 snRNP-independent sites), and 1,044 sites were specific to U1-70K KD (SRSF1-independent sites). However, 222 sites were significantly affected by both KDs, either in the same or opposite direction (referred to here as “shared PASs”) (**Fig. 2C**); this overlap was significantly higher than predicted by chance (hypergeometric test p-value = 9.78e-22). We identified 87 shared PASs with effects in the same direction, and 135 shared PASs with effects in the opposite direction. For example, in the gene *TMEM242*, both SRSF1 and U1-70K KD increased usage of a particular PAS (**Fig. 2D**). In contrast, in *GPR107*, SRSF1 and U1-70K KDs had opposite effects on the usage of a particular PAS, with SRSF1 KD leading to decreased usage and U1-70K KD leading to increased usage (**Fig. 2E**). That SRSF1 and U1-70K KDs can shift shared PASs in the same or opposite directions suggests that, rather than acting redundantly, the two factors play distinct regulatory roles in PAS selection at shared sites, analogous to their relationship in splicing regulation, where SRSF1 can either promote or inhibit exon recognition by U1 snRNP and the core spliceosome^44,45^.

**Figure 2.**
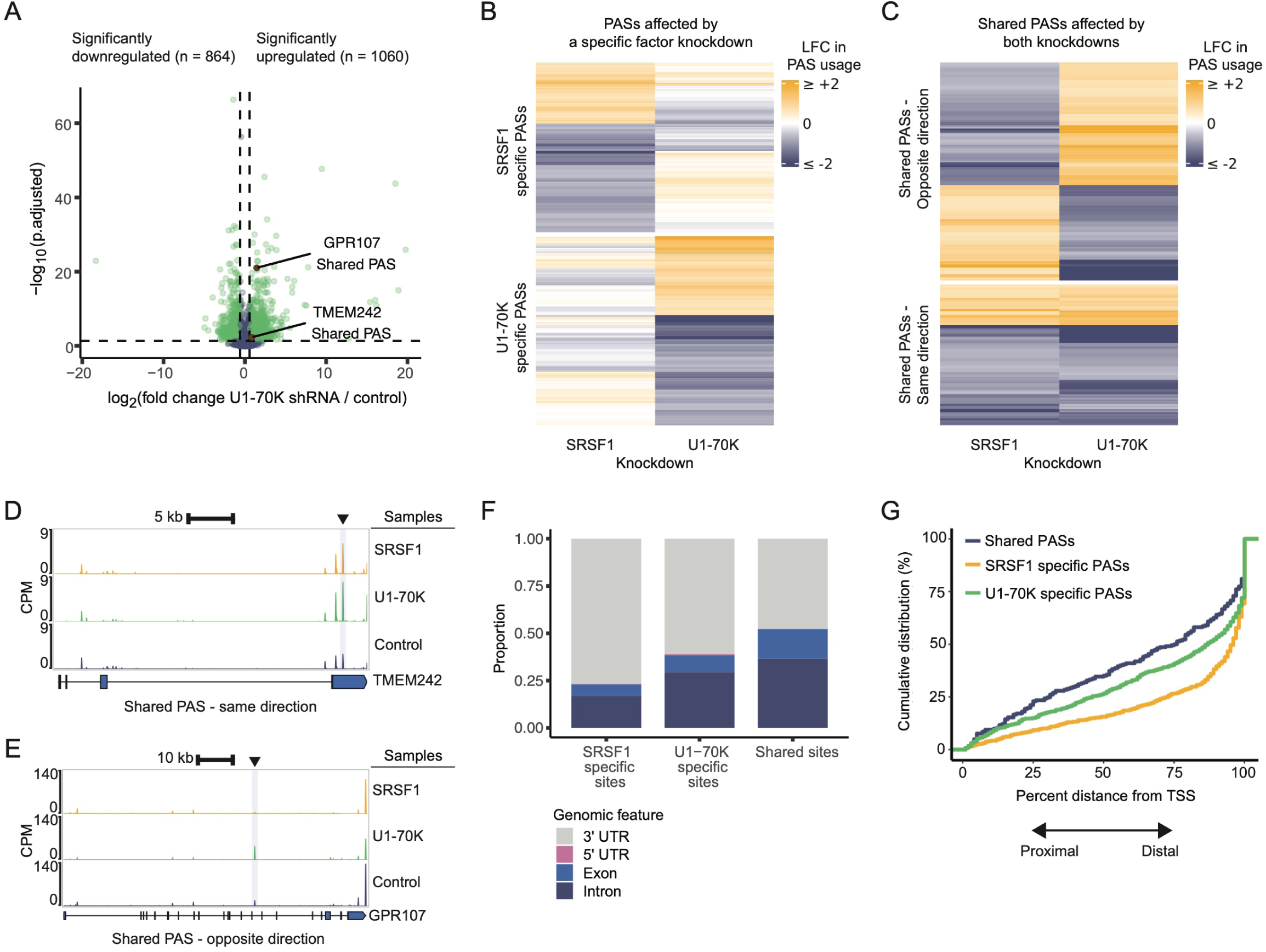
SRSF1 and U1 snRNP regulate a shared set of PASs. A) Volcano plot of changes in relative PAS usage upon U1-70K KD, as calculated by DEXseq (n = 3). Each dot is a PAS, and green dots are significantly changed PASs while purple are unchanged. Brown dots represent particular examples shown later in the figure. 1060 PASs had significantly upregulated usage, 864 sites had significantly downregulated usage, and 14,617 sites were unaffected. Significantly differentially used PASs had p.adjusted < 0.05, abs(log_2_(fold change)) > log_2_(1.5). B) Heatmap of the log_2_ fold changes (LFC) in significantly affected PASs upon each KD (n = 937 SRSF1 specific PASs, n = 1044 U1-70K specific PASs). Heatmap scale fixed from −2 to 2. Changes in PAS usage were clustered by Euclidean distance and Ward.D2 linkage using ComplexHeatmap. C) Heatmap of the log_2_ fold changes (LFC) in significantly affected PASs upon both KDs (n = 87 shared PASs in the same direction, n = 135 shared PASs in the opposite direction). Heatmap scale fixed from −2 to 2. Changes in PAS usage were clustered by Euclidean distance and Ward.D2 linkage using ComplexHeatmap. D-E) Genome browser tracks for 3′-end sequencing data for the genes *TMEM242* (D) and *GPR107* (E). F) Proportions of SRSF1 specific, U1-70K specific, and shared PASs with significantly altered usage (p.adjusted < 0.05, abs(log_2_(fold change)) > log_2_(1.5)) upon the respective factor KD mapped to genomic features. Shared PASs changing either in the same or opposite directions upon the KDs were both analyzed. G) Empirical cumulative distributions of the relative positions of significantly altered PASs across genes between the TSS and the most distal PAS detected in our 3′-end sequencing experiments. Shared PASs changing either in the same or opposite directions upon the KDs were both analyzed.

We next cataloged where the SRSF1 and U1 snRNP shared PASs reside within genes. Over a third of shared PASs were intronic (**Fig. 2F**), consistent with the established role of U1 snRNP in regulating intronic polyadenylation^36,37^. Notably, a higher proportion of shared PASs were intronic than were for PASs specifically affected by U1-70K KD alone (**Fig. 2F**). Moreover, rather than having overlapping distributions, shared PASs were positioned more proximal to the TSS compared to PASs specifically affected by SRSF1 or U1-70K KD (**Fig. 2G**). Together, these patterns suggest that SRSF1’s influence on PAS usage varies across genes, with some sites predominantly sensitive to SRSF1 and enriched at distal positions within genes, and others sensitive to both SRSF1 and U1 snRNP and enriched more proximally throughout gene bodies.

#### SRSF1 binds towards the start of 3′ UTRs, near SRSF1-sensitive PASs

Given that PASs predominantly regulated by SRSF1 are enriched in 3′ UTRs (**Fig. 1G**), we next aimed to determine how SRSF1 shapes PAS usage at U1 snRNP-independent sites within 3′ UTRs. Consistent with our aggregated analyses of SRSF1-sensitive PASs across genomic features (**Fig. 1H**), inspection of PAS usage across 3′ UTRs revealed a shift toward the use of more distal sites compared to proximal sites upon SRSF1 KD (**Fig. 3A, S3A**). These proximal and distal sites could be located in the same or different last exons. To independently assess this shift using an orthogonal approach, we applied the hybrid-internal-terminal (HIT) index pipeline, which identifies the inclusion rates of alternative last exons from whole cell RNA-seq data^46^. Consistent with our 3′-end sequencing results, this analysis revealed that SRSF1 KD shifts last exon usage distally, further supporting a role for SRSF1 in promoting proximal PAS selection (**Fig. 3B, S3B**).

**Figure 3.**
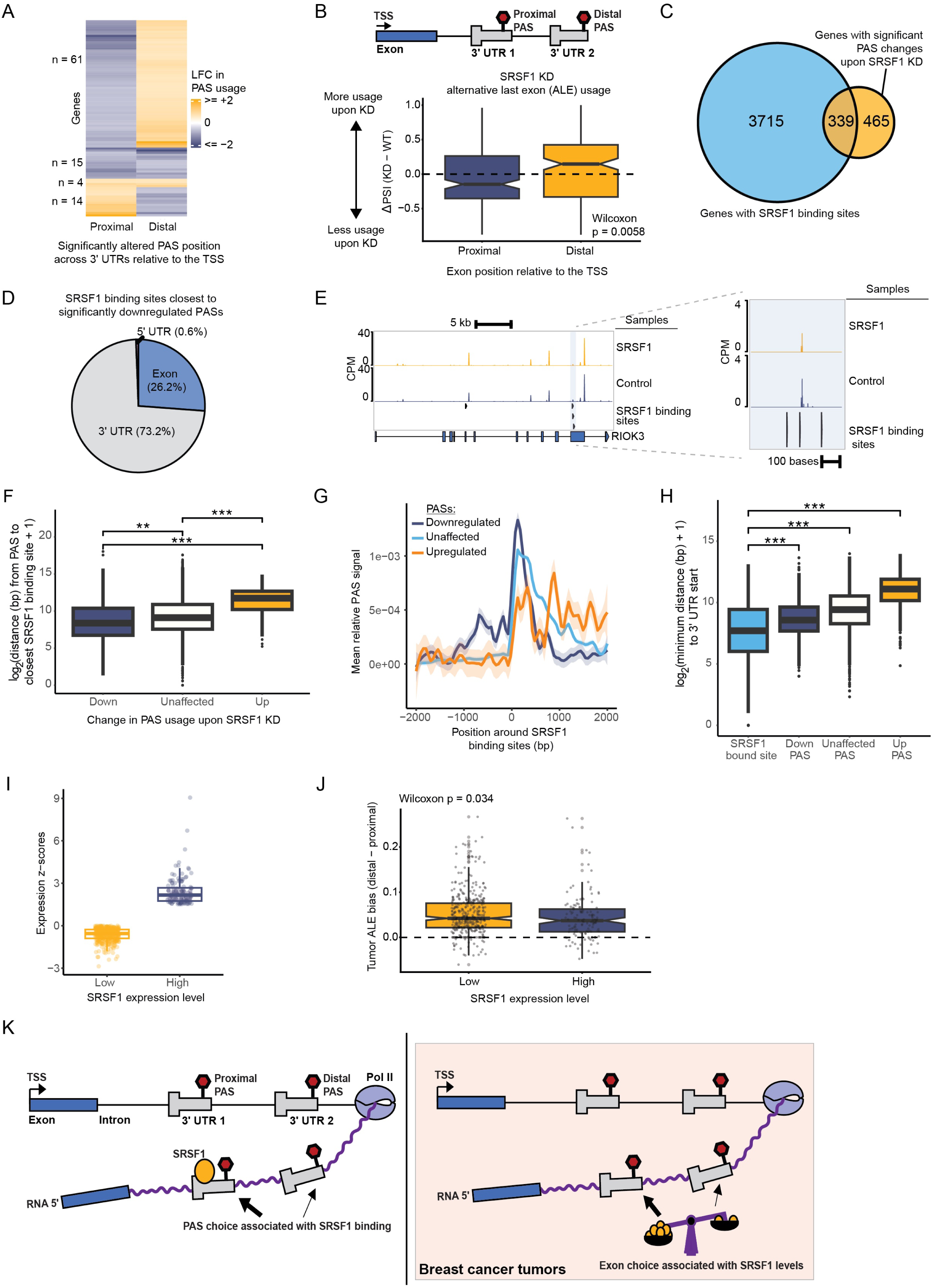
SRSF1 binds towards the start of 3′ UTRs, near SRSF1-sensitive PASs. A) Heatmap of the log_2_ fold change (LFC) in PAS usage across one or more 3′ UTRs with two significantly altered proximal/distal PASs (n = 94). Changes in PAS usage were clustered by Euclidean distance and Ward.D linkage using ComplexHeatmap; the number of genes per clustered pattern in changed PAS usage is indicated to the left of the heatmap. B) (Top) Schematic of an example gene with proximal/distal PASs on alternative last exons (3′ UTRs). (Bottom) HITindex pipeline analyses of the delta percent spliced in (PSI) of proximal and distal alternative last exons (ALEs) in whole cell RNA upon SRSF1 KD. The analysis was run on exons in genes (n = 207 genes) with significant shifts in the delta PSI (|delta PSI| > 0.1, FDR < 0.05). C) Venn diagram of genes containing SRSF1 binding sites identified in published PIE-seq data (Ruan *et al.* 2023) and genes with significantly altered PAS usage following SRSF1 KD. D) Genomic feature mapping for the PIE-seq SRSF1 binding sites nearest to PASs significantly downregulated upon SRSF1 KD (n = 168 binding sites). E) Genome browser tracks of our 3′-end sequencing data and PIE-seq SRSF1 binding data for *RIOK3*. F) Boxplots of the log_2_ distance in basepairs (bp) (+1 bp pseudocount) between each PAS and its closest PIE-seq SRSF1 binding site. Only distances between PASs and binding sites that both mapped to the 3′ UTR were analyzed. (Downregulated PASs n = 164, upregulated PASs n = 94, unaffected PASs n = 3018). G) Metagene plot of PAS signal from control samples (n = 3) surrounding 3′ UTR-mapped PIE-seq SRSF1 binding sites. PAS signal was subsetted based on the effect of SRSF1 KD. Within each subset, to remove noise, we filtered out binding sites surrounded by the bottom 25% lowest total PAS signal. Within +/-2 kb of binding sites, the relative PAS usage was averaged across the binding sites profiled. H) 3′ UTRs containing both PIE-seq SRSF1 binding sites and PASs or only PASs were assessed for the log_2_ distances (in basepairs, +1 bp pseudocount) between the start of each 3′ UTR and the nearest binding site or PAS, with PASs subsetted by the effect of SRSF1 KD. Number of sites analyzed and median nearest distances: 1012 bound sites, 208 bp; 445 downregulated sites, 382 bp; 3993 unaffected sites, 686 bp; 281 upregulated sites, 2190 bp. I) SRSF1 mRNA expression z-scores relative to diploid samples across breast tumor samples, as assigned by cBioPortal. High SRSF1-expressing tumors had z-scores > 1.5. Low SRSF1-expressing tumors had z-scores < 0. J) The overall median ALE bias (distal - proximal PAS usage) for each tumor was calculated using the HITindex pipeline for TCGA breast cancer samples with varying levels of SRSF1 expression. High SRSF1-expressing tumors had an expression z-score > 1.5. Low SRSF1-expressing tumors had a z-score < 0. K) (Left) Schematic of SRSF1 binding near proximal PASs in 3′ UTRs, promoting their usage. Loss of SRSF1 leads to PAS site choice with decreased selection of the proximal PAS. (Right) Promotion of proximal PAS usage by SRSF1 is recapitulated in breast tumor samples with ALE usage and differential SRSF1 levels.

To investigate whether SRSF1 binding could directly influence PAS usage, we analyzed genes that contained SRSF1-sensitive PASs and overlapped them with genes containing previously reported SRSF1 binding sites^47^ (n = 339) (**Fig. 3C**). We observed that the SRSF1 binding sites closest to SRSF1-sensitive, downregulated PASs were mostly in 3′ UTRs (∼73% of sites) (**Fig. 3D, S3C).** Using 3′ RACE, we validated one such site in *RIOK3*, which has a proximal PAS in the 3′ UTR, near SRSF1 binding sites, which is downregulated upon SRSF1 KD (**Fig. 3E, S3D**). Indeed, SRSF1-sensitive, downregulated PASs were located closer to SRSF1 binding sites and showed greater enrichment around them than other PASs in 3′ UTRs (**Fig. 3F-3G, S3E-S3G**). This spatial relationship was not observed around other RNA binding proteins that bind within 3’ UTRs, including NOVA1 and PUM2 (**Fig. S3H-S3I**). On average, the most upstream SRSF1 binding sites within 3′ UTRs were positioned closest to the start of 3′ UTRs, followed by SRSF1-sensitive, downregulated PASs, and then by unaffected and upregulated PASs further downstream (**Fig. 3H, S3J**). This spatial ordering – with bound SRSF1 upstream of SRSF1-sensitive proximal PASs – is consistent with a model in which SRSF1 binding near the start of the 3′ UTR favors the usage of nearby PASs (**Fig. 3K**, **left**).

#### SRSF1 expression levels in breast cancer tumors are associated with alternative last exon usage

SRSF1 is often amplified and upregulated in breast cancer^48,49^, where it promotes oncogenic splicing events^50^. Therefore, we asked whether altered SRSF1 levels are associated with shifted 3′-end selection within a tumor context. To address this question, we retrieved patient sample data from The Cancer Genome Atlas (TCGA)^51^ and analyzed human breast cancer tumors previously characterized to express either high (n = 128) or low (n = 403) levels of SRSF1^48,52^ **(Fig. 3I)**. Using TCGA primary tumor whole cell RNA-seq data, we analyzed the data for changes in last exon usage using the HITindex pipeline^46^. Despite the heterogeneity inherent to primary tumor samples, we saw that tumors with low SRSF1 levels had significantly distally shifted last exon usage compared to tumors with high SRSF1 levels (**Fig. 3J** and **3K**, **right**), consistent with our observations of distally shifted PAS usage in the absence of SRSF1 in HEK293T cells (**Fig. 3B, S3B**). Therefore, the shift in last exon usage we observe in HEK293T cells is recapitulated in breast tumors with differing SRSF1 levels, suggesting that SRSF1’s effect on PAS selection extends to disease-relevant contexts. These results suggest that, in addition to splicing changes, SRSF1-driven changes in exon usage have the potential to contribute to oncogenic isoform production in breast cancer.

#### Interactions between SRSF1 and Pol II are dependent on U1 snRNP

In contrast to the uniquely SRSF1-sensitive PASs, the U1/SRSF1 shared PASs are far more proximal to TSSs (**Fig. 2G)**. As U1 snRNP modulates PAS usage by stimulating Pol II elongation^33,34^, and SR proteins can act co-transcriptionally to shape RNA processing^15^, we hypothesized that physical interactions between U1 snRNP, SRSF1, and Pol II shape co-transcriptional PAS regulation at shared PASs. Consistent with our hypothesis, we found that AlphaFold3 modeling^53^ predicted an interaction interface between U1-70K and phosphorylated SRSF1 (**Fig. 4A, S4A-S4B**). Importantly, mapping the predicted interacting domains of SRSF1 and U1-70K onto a cryoEM structure of the transcribing Pol II - U1 snRNP complex^32^ did not reveal steric clashes between SRSF1 and the transcription elongation complex (**Fig. 4B**).

**Figure 4.**
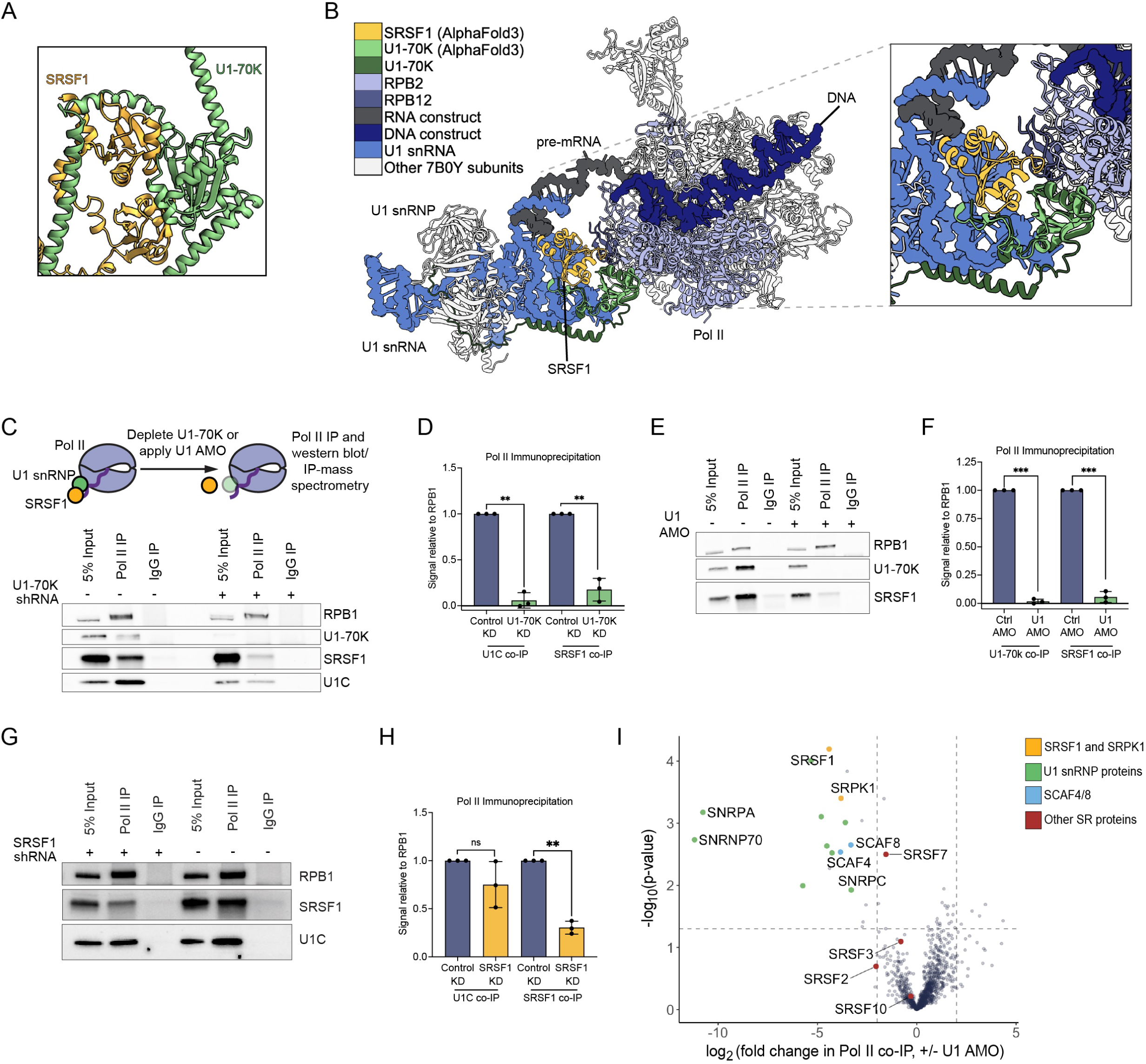
Interactions between SRSF1 and Pol II are dependent on U1 snRNP. A) Zoom-in view of the folded and interacting domains in the AlphaFold3 (AF3) predicted structure of full-length phosphorylated SRSF1 and U1-70K. 12 serines in SRSF1 were modeled with phosphorylation from residue 205 to 227. Structure is colored by protein identity. Interface predicted template modeling (ipTM) score = 0.31; fraction disordered = 0.24. B) AF3 structure prediction of interacting phosphorylated SRSF1 and U1-70K domains mapped onto the structure of a transcribing Pol II-U1 snRNP complex (PDB: 7B0Y from Zhang *et al.* 2021). Subunits from 7B0Y unless otherwise indicated in the key. The following sequences are shown: SRSF1 residues 5-105 from the AF3 prediction, U1-70K residues 100-205 from the AF3 prediction, U1-70K residues 27-205 from 7B0Y, and all other proteins and nucleic acids from 7B0Y in full. Inset shows a zoomed-in view of the Pol II-U1-70K-SRSF1 interaction interface. C) (Top) Schematic of the experimental setup in which Pol II-U1 snRNP interactions are disrupted using a U1-70K-targeting shRNA or U1 AMO, followed by Pol II immunoprecipitation (IP), western blotting, and/or IP-mass spectrometry (IP-MS). (Bottom) Pol II IP performed in the presence of a U1-70K-targeting shRNA (+) or non-silencing shRNA (-). D) Quantification of C. E) Pol II IP performed in the presence of U1 AMO (+) or a scrambled control AMO (-). F) Quantification of E. G) Pol II IP performed in the presence of an SRSF1-targeting shRNA (+) or non-silencing shRNA (-). H) Quantification of G. I) Analysis of IP-MS data for Pol II IP in the presence of the U1 AMO compared to a scrambled control AMO (n = 4). Significance threshold set at |log_2_ fold change in Pol II co-IP| ≥2 and −log_10_ p-value ≥1.3.

To test our hypothesis experimentally, we asked whether SRSF1 associates with Pol II by co-immunoprecipitation (co-IP) from HEK293T nuclear lysates, and whether this interaction depends on U1 snRNP. To disrupt Pol II-U1 snRNP interactions, we used two approaches: a U1-70K-targeting shRNA to deplete U1-70K protein, and a U1 snRNA-targeting AMO (U1 AMO) to disrupt U1 snRNP interactions with nascent RNA and Pol II^29,54^ (**Fig. 4C, top**). In our control Pol II co-IPs, we observed a strong association between Pol II, SRSF1, and U1 snRNP, as reported previously^15^ (**Fig. 4C, bottom and 4E**). Upon applying either perturbation, interactions between U1 snRNP subunits and Pol II decreased, as expected (**Fig. 4C-4F**). Notably, co-IP between SRSF1 and Pol II was also significantly reduced (**Fig. 4C-4F**), suggesting that Pol II-SRSF1 interactions rely on U1 snRNP. Conversely, upon shRNA KD of SRSF1, U1 snRNP largely remained associated with Pol II (**Fig. 4G-4H**), suggesting SRSF1 does not reciprocally influence interactions between Pol II and U1 snRNP. Together, these results demonstrate that SRSF1 interacts with Pol II in a U1 snRNP-dependent manner.

We next asked whether U1 snRNP mediates Pol II interactions for a broader set of factors beyond SRSF1. To address this, we performed IP-mass spectrometry on Pol II immunoprecipitated from cells treated with a U1 or scrambled AMO. We identified expected members of the Pol II interactome in the scrambled AMO control IPs, including elongation factors (**Fig. S4C**), and proteins functionally enriched for RNA polymerase II activity, pre-mRNA intronic binding, and spliceosome components (**Fig. S4D-S4E**). As expected, U1 AMO treatment led to the loss of U1 snRNP subunits and SRSF1 from Pol II, confirming our western blot results (**Fig. 4I**). Notably, beyond U1 snRNP subunits and SRSF1, only a small number of additional proteins showed significantly decreased Pol II interactions upon U1 AMO treatment (**Table S1**), indicating that U1 snRNP mediates associations between Pol II and a specific, limited set of factors rather than broadly remodeling the Pol II interactome. Among these, U1 AMO treatment notably decreased Pol II interactions with SRPK1, a kinase known to target SR dipeptide motifs^55^, and with SCAF4 and SCAF8, which have previously been shown to regulate the Pol II elongation rate, transcription termination, and PAS selection (**Fig. 4I**)^25^. Consistent with reports that other SR proteins bind to Pol II but not U1 snRNP^42^, other Pol II-interacting SR proteins were not significantly affected by the U1 AMO treatment (**Fig. 4I**). These results demonstrate that U1 snRNP is required for a small but functionally coherent set of factors to maintain robust interactions with Pol II.

#### SRSF1 slows Pol II elongation and limits transcription readthrough

Given that U1 snRNP accelerates Pol II elongation to influence PAS usage^33^, and that SRSF1 depends on U1 snRNP for Pol II association (**Fig. 4**), we hypothesized that SRSF1 could similarly shape PAS choice by altering transcription dynamics. We therefore sought to determine whether SRSF1 modifies Pol II behavior. To this end, we first looked at the effects of SRSF1 on transcription and Pol II density. In SRSF1 knockdown cells, we performed transient transcriptome sequencing (TT-seq) to measure nascent transcription and precision run-on sequencing (PRO-seq) to map the position of Pol II along genes (**Fig. 5A, S5A-S5F**). SRSF1 KD decreased nascent transcription of 3,331 genes (**Fig. S5G**), which correlated well with our PRO-seq data (R = 0.79; **Fig. S5H-S5I**). Nascent transcription was affected evenly across genes, ruling out gene body premature transcription termination as responsible for the decreased transcription (**Fig. S5J**). Consistent with these results, we observed lower protein levels of Pol II expression (total, Ser2P, Ser5P) (**Fig. S5K-S5M**).

**Figure 5.**
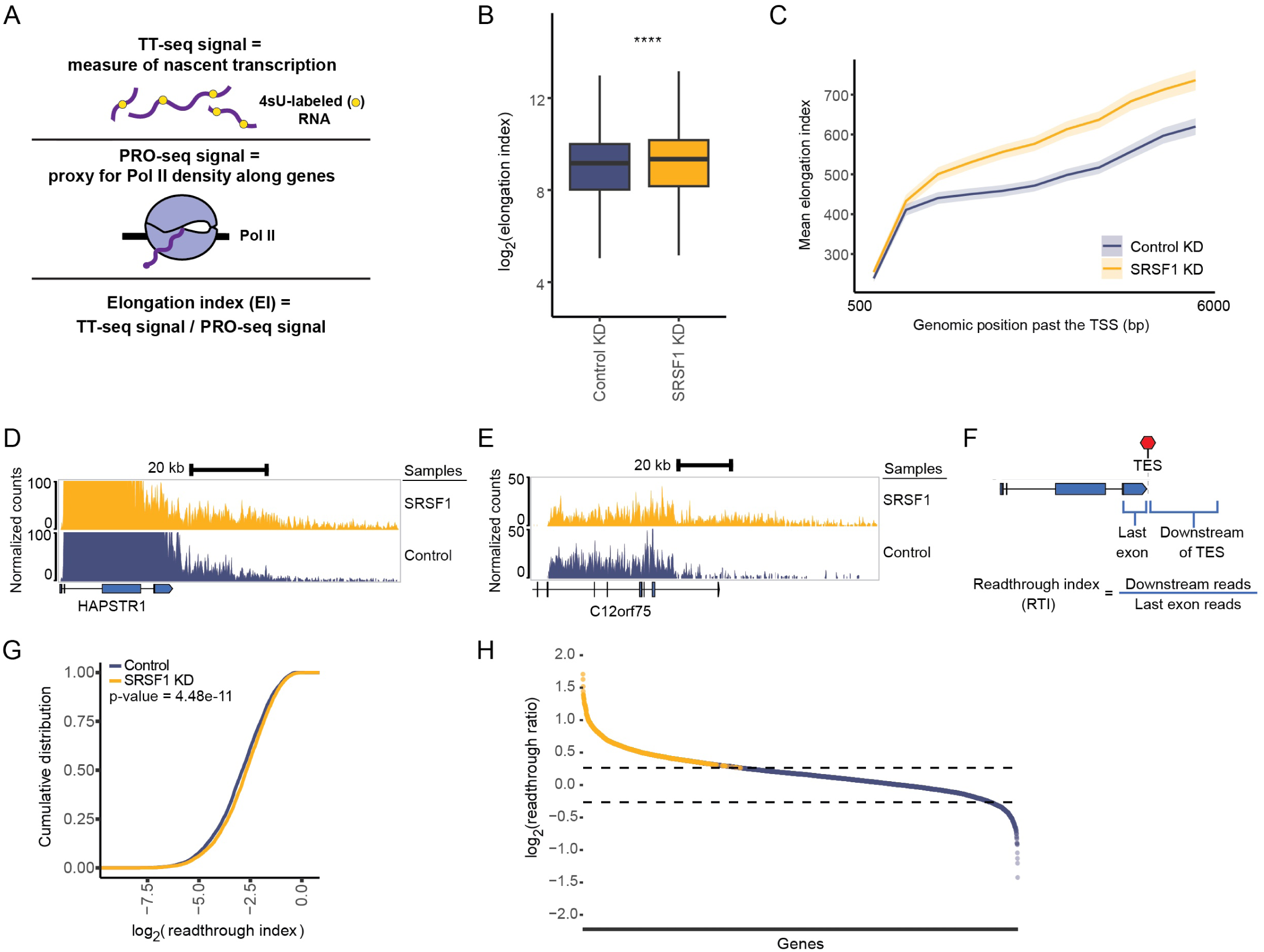
SRSF1 slows Pol II elongation and limits transcription readthrough. A) Schematic: the ratio of TT-seq and PRO-seq data was used to calculate the elongation index (EI). Nascent RNA was labeled with 4-thiouridine (4sU) for TT-seq. B) Boxplots of the log_2_(EI) plotted by KD condition. We evaluated the EI up to the first 50.5 kb of each gene, with the TSS and first 500 bp removed from the analysis. The 50 kb region was binned in 500 bp bins, and bins with the lowest 5% of counts were dropped. Genes were filtered for those with unaffected nascent transcription, as determined by DESeq2 analysis of TT-seq data (p.adjusted > 0.05, abs(log_2_(fold change)) < 1.5). Outlier points not shown. Significance determined by Wilcoxon test. C) Metagene plot of the mean EI per bin (+/- 95% confidence interval) from 500 bp to 6 kb past the dominant TSS. Bins used were 500 bp. Bins with the lowest 5% of counts were dropped. The TSS and first 500 bp downstream were removed from the analysis. Genes were filtered for those with unaffected nascent transcription, as determined by DESeq2 analysis of TT-seq data (p.adjusted > 0.05, abs(log_2_(fold change)) < 1.5). D-E) Examples of increased readthrough in the regions downstream of the genes *HAPSTR1* (D) and *C12orf75* (E) upon SRSF1 KD. Reads were normalized by size factors derived from *Drosophila* spike-ins using DESeq2. F) For each gene, the readthrough index (RTI) is the ratio of the average, length and spike-in normalized TT-seq value of the region 20 kb downstream of the TES compared to the average, length and spike-normalized TT-seq value of the last exon containing the TES. G) Cumulative distribution of the RTI in both control and SRSF1 KD conditions. P-value determined by the Kolmogorov–Smirnov test. H) Log_2_ readthrough ratios (i.e. log_2_(RTI for SRSF1 KD / RTI for control KD)) for each gene, ordered from largest to smallest. Genes where all replicate ratios (n = 3) and the averaged ratio are ≥1.2 are determined to have reproducible readthrough events (n = 880, colored in yellow).

We next calculated the effect of SRSF1 on the Pol II elongation rate. Relative rates of transcription elongation can be estimated by the elongation index (EI), the ratio of TT-seq signal to PRO-seq signal along genes, which captures the density of nascent transcripts relative to the density of Pol II (**Fig. 5A**). Loss of SRSF1 led to an increase in the elongation index across gene bodies (**Fig. 5B-5C**). Thus, in normal conditions, SRSF1 acts to reduce the average Pol II elongation index across genes, in contrast to U1 snRNP, which accelerates elongation^33^.

Previous studies have established that Pol II elongation and termination kinetically compete with one another^24,25,27^. Because SRSF1 reduces the Pol II elongation index under normal conditions, we predicted that SRSF1 depletion should lead to increased transcription readthrough as a consequence of less efficient termination. Indeed, using our TT-seq data, we observed many genes exhibiting run-on transcription, including *HAPSTR1* and *C12orf75* (**Fig. 5D-5E**). To quantify this effect globally, we calculated a readthrough index (RTI), comparing transcription 20 kb downstream of the dominant transcription end site (TES) to transcription in the last exon (**Fig. 5F**). We found a significant increase in transcription readthrough upon SRSF1 KD (**Fig. 5G**), driven by 880 genes displaying reproducible differences in downstream transcription (**Fig. 5H**). Together with our EI analysis, these data support a model in which SRSF1 shapes Pol II dynamics, influencing Pol II elongation as well as termination.

## Discussion

In this study, we demonstrate that SRSF1 plays a critical role in regulating PAS usage through two complementary mechanisms whose relative importance varies across genomic contexts: direct RNA binding nearby PASs within the 3′ UTR, where SRSF1’s effects are largely independent of U1 snRNP, and co-transcriptional coordination with U1 snRNP at sites distributed throughout the gene body (**Fig. 6A-6B**). For sites predominantly sensitive to SRSF1, the proximity between SRSF1 binding sites and SRSF1-sensitive PASs near the start of the 3′ UTR suggests a model where bound SRSF1 promotes the usage of nearby proximal PASs. When SRSF1 is not present, PAS choice may occur over a wider radius, shifting usage distally within the 3′ UTR (**Fig. 3K, left**).

**Figure 6.**
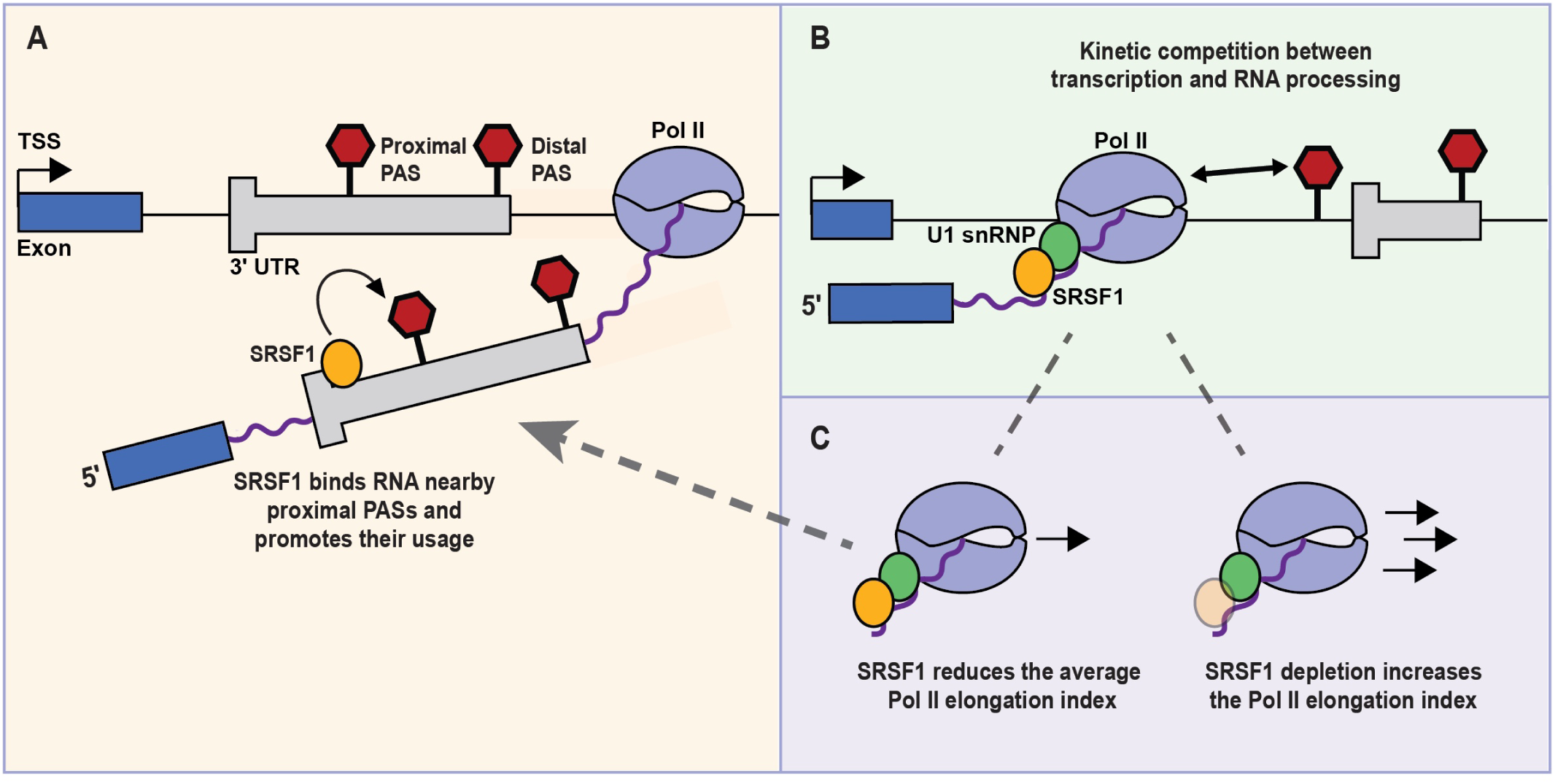
Model for how SRSF1 regulates 3′-end selection through complementary mechanisms with differential dependence on U1 snRNP. A) At PASs where SRSF1’s effects are largely U1 snRNP-independent, SRSF1 binds nearby proximal sites in the 3′ UTR to promote their usage. B) At PASs sensitive to both SRSF1 and U1 snRNP, SRSF1 tunes the elongation rate of Pol II. Consistent with the kinetic model of PAS regulation, Pol II speed influences the likelihood of PAS selection and transcription termination at different sites. C) The mechanisms proposed in A and B are not mutually exclusive and likely operate in tandem at individual PASs.

How SRSF1 shapes proximal PAS usage within the 3′ UTR remains to be fully determined. If SRSF1 binding acts directly at these sites, several mechanisms could account for this relationship. One possibility is that SRSF1 recruits CPA machinery to nearby PASs. This model is consistent with studies showing that SR proteins, including SRSF1, recruit CPA factors in a viral context^56–59^, and that arginine/serine-rich domains can mediate interactions between splicing factors and CPA components^60,61^. Alternatively, SRSF1 may compete with RNA binding proteins that inhibit CPA at these sites. In either case, the involvement of a splicing factor in PAS regulation is consistent with the long-established finding that splicing and CPA are mutually stimulatory^62,63^, and raises the possibility that the positioning of SRSF1 binding sites towards the beginning of the 3′ UTR functions to couple splicing, cleavage, and PAS choice.

SRSF1 is frequently upregulated in cancer, where it promotes oncogenic splicing events^48–50,64^. Our finding that breast cancer tumors with differing SRSF1 levels exhibit corresponding shifts in last exon usage (**Fig. 3J**) suggests that SRSF1-driven changes in PAS selection also contribute to oncogenic isoform production. As intronic polyadenylation events can inactivate tumor suppressor genes^9^, it will be important to assess if SRSF1 overexpression is sufficient to induce oncogenic alternative polyadenylation events and to identify novel pathways through which SRSF1 levels influence cancer phenotypes.

Beyond PASs where SRSF1’s effects predominate, SRSF1 also converges with U1 snRNP on a set of PASs distributed throughout the gene body that are sensitive to both factors (**Fig. 2**). That the two KDs shift these shared PASs in the same or opposite directions suggests that, rather than acting as a single entity, their relationship in PAS selection is analogous to their relationship in splicing regulation, where U1 snRNP serves as a core regulator and SRSF1 either promotes or inhibits its effects. One paradigm for how these factors shape shared PAS usage is the “window of opportunity” model, where the Pol II elongation rate influences the likelihood of PAS selection and termination at different sites, with slower elongation rates favoring recognition of proximal sites and faster rates favoring distal sites^24–28,65–69^. Within this framework, we demonstrate that SRSF1 reduces the average Pol II elongation index (**Fig. 5**), while U1 snRNP has been shown to accelerate Pol II^33^, suggesting that these factors tune the final Pol II elongation rate for optimal PAS selection.

Notably, modulating transcription dynamics may reinforce SRSF1’s direct RNA binding effects even at predominantly SRSF1-sensitive PASs (**Fig. 6C**), as the effect of SRSF1 on elongation is compatible with its impact on PAS usage in the 3’ UTR (**Fig. 3**). Namely, under normal conditions, SRSF1-mediated slowing of Pol II may provide additional time for CPA machinery to recognize proximal PASs, reinforcing the promotion of proximal PAS usage by RNA-bound SRSF1. Upon SRSF1 depletion, the combined loss of RNA binding and increased Pol II elongation rate would both favor distal PAS selection, suggesting that these two mechanisms act in a complementary manner to ensure robust proximal PAS usage. Consistent with this model, faster endogenous Pol II elongation rates are associated with increased downstream PAS usage in human cells^28^.

A physical basis for coordination between SRSF1 and U1 snRNP in PAS regulation is supported by our biochemical and proteomic data (**Fig. 4**). Past studies have used the U1 AMO to disrupt Pol II - U1 snRNP interactions and characterize PAS regulation by U1 snRNP^29,30,33,34^. Building upon this work, we demonstrate that the same perturbation prevents a small group of additional factors — notably SRSF1, SCAF4, and SCAF8 – from interacting with Pol II. These three factors have now all been shown to influence Pol II speed, transcription termination, and PAS selection^25^. Given their physical dependence on U1 snRNP for Pol II association, we posit that some phenotypes attributed to U1 snRNP upon U1 AMO application may instead be mediated by disrupted SRSF1, SCAF4, or SCAF8 interactions with Pol II. It is also plausible that U1 snRNP works in coordination with these factors at the interface between Pol II dynamics and PAS selection.

Our findings fit nicely into the recently proposed “U1 relay model,” which suggests that mutually exclusive interactions between U1 snRNP, SPT6, and CPA machinery and their resulting orientations on Pol II relative to nascent RNA are important for shaping 3′-end processing^70^. Within this framework, individual U1 snRNP complexes cycle on and off Pol II during splicing. Our observation that SRSF1, SCAF4, and SCAF8 depend on U1 snRNP for Pol II association implies that these factors may cycle alongside U1 snRNP. Therefore, we propose that these proteins are strong candidates to directly contact elongation machinery and influence co-transcriptional interactions among proteins bound to Pol II.

Overall, we demonstrate that SRSF1 plays an important role in shaping PAS selection with site-specific differential degrees of independence from U1 snRNP. By acting through direct RNA binding and U1 snRNP-mediated Pol II interactions, SRSF1 bridges splicing, transcription dynamics, and PAS selection in a manner that expands our understanding of how pre-mRNA processing factors coordinate multiple layers of gene regulation^25,33–39,71–73^.

## METHODS

### EXPERIMENTAL MODEL DETAILS

#### Cell culture

HEK293T cells were used for all experiments. They were cultured at 37°C and 5% CO_2_ in DMEM media (Gibco #11995-065) supplemented with 10% FBS and 1% penicillin/streptomycin. Cells were checked routinely for absence of mycoplasma contamination using the ATCC Universal Mycoplasma Detection Kit (#30-1012K).

#### Inducible shRNA cell line generation

Inducible shRNA cell lines were generated using the Horizon Discovery TRIPZ Inducible Lentiviral shRNA system. TRIPZ plasmids used include the non-silencing negative shRNA control (#RHS4743), SRSF1-targeting shRNA (#RHS4696-200705213, clone #V2THS_149883), and SNRNP70-targeting shRNA (#RHS4696-200772835, clone #V3THS_357957). To generate virus for inducible shRNA lines, approximately 0.3-0.5 million HEK293T cells were seeded on day 0. On day 1, TRIPZ plasmids were packaged along with the psPAX2 and pMD2.G lentiviral packaging plasmids from Addgene (#12260 and #12259 respectively) using the Lipofectamine 3000 Transfection Reagent as instructed by the manufacturer (Thermo #L3000001). Virus was collected on day 3. Collected media was filtered through a low protein binding 0.45 µM filter, and 1.5 ml of the filtered virus was added onto approximately 0.5 million HEK293T cells. One negative control well containing HEK293T cells did not receive any virus. On day 4, the media was replaced with fresh DMEM media. On day 5, cells were split to 33% confluency and puromycin was added to 2 µg/ml. Every following two days, fresh DMEM media was added to cells along with fresh puromycin, and this was repeated until no live cells remained in the negative control well. The resulting transduced cells were used for subsequent experiments. Successful knockdown of SRSF1 or U1-70K upon doxycycline-induction of these cells was confirmed by western blot. The sequences of TRIPZ shRNA constructs in the generated cell lines were also confirmed through Sanger sequencing.

### METHOD DETAILS

#### siRNA knockdown

The following siRNAs were purchased from Horizon Discovery: a SRSF1-targeting siRNA (J-018672-09-0005), a U1-70K-targeting siRNA (custom SNRNP70 duplex CTM-739231, sense 5’ A.A.G.G.G.U.A.G.G.U.G.U.C.U.C.A.U.U.U.U.U 3’ and antisense 5’ 5’-P.A.A.A.U.G.A.G.A.C.A.C.C.U.A.C.C.C.U.U.U.U 3’), and a non-targeting siRNA control (#1 D-001810-01-05). Approximately 1-2 million HEK293T cells were seeded on day 0. On day 1, siRNAs were added to cells at 25 nM using the Lipofectamine RNAiMAX Transfection Reagent (Invitrogen #13778075) as instructed by the manufacturer. On day 3, approximately 2.5 million cells were seeded into experimental and reference plates for collection on day 4. Whole cell RNA and protein were collected from reference plates, and knockdown of SRSF1 or U1-70K was confirmed by western blot.

#### Western blot analysis

The following whole cell and fractionated samples were used for western blotting: U1-70K and SRSF1 shRNA KD samples from 3′-end sequencing experiments, U1-70K and SRSF1 shRNA KD samples from IP experiments, U1 AMO samples from IP experiments, SRSF1 siRNA KD samples from TT-seq experiments, and SRSF1 siRNA KD from PRO-seq experiments, along with their appropriate controls. Total protein samples were normalized prior to performing western blot either with a Qubit protein assay (ThermoFisher Q33212) or a BCA assay (Pierce #23227). SDS or LDS buffer (Invitrogen NP0007) was then added to 1X concentration, with 2-mercaptoethanol (Sigma-Aldrich M6250) added at a final concentration of 2.5%. Samples from immunoprecipitation (IP) were prepared as detailed in the section on IP experiments.

Prior to western blotting, samples were boiled for 5 minutes and loaded onto a NuPAGE 4-12% Bis-Tris gradient gel (ThermoFisher). For IP samples, 5% input was loaded. Samples were separated for 1 hour at 160 V in 1X MOPS buffer (ThermoFisher NP0001) and transferred onto a nitrocellulose membrane in an electrotransfer chamber for 2 hours at 120 V in standard transfer buffer. Transfer was assessed by Ponceau stain. Samples were blocked in 5% milk in 1X TBST for 1 hour at room temperature. The membranes were incubated at 4°C on a shaker, in primary antibody and 5% milk overnight, with primaries indicated in the table below. The next day, blots were washed four times in 1X TBST for 5 minutes before incubating with a secondary antibody (listed in table below) and 5% milk at room temperature for 1 hour. Membranes were subsequently washed twice more with 1X TBST for 15 minutes and 1X PBS once for 5 minutes before imaging on the Bio-Rad ChemiDoc Imaging System.

Quantification of western blot results was performed using Image Lab Software volume tools, using the adjusted volume (intensity) for each band. For quantification of the IP experiments, the SRSF1 to RPB1 signal ratio (and U1 to RPB1 signal ratio) was first determined for each condition. Next, the relative intensity was calculated as experimental condition/control condition for both SRSF1 and U1. The 5% input lane was not used in the quantification. For quantification of Pol II levels (including total RPB1, Ser2, and Ser5), and quantification of SRSF1 and U1-70k knockdown in PRO-seq samples, the signal of the protein of interest was calculated and normalized to the loading control (GAPDH or beta-tubulin), and compared between control and experimental conditions. For all quantifications, a paired t-test was used to compare conditions within replicates. * indicates p <0.05, ** indicates p < 0.01, *** indicates p < 0.001, and **** indicates p < 0.0001.

Table of antibodies used:

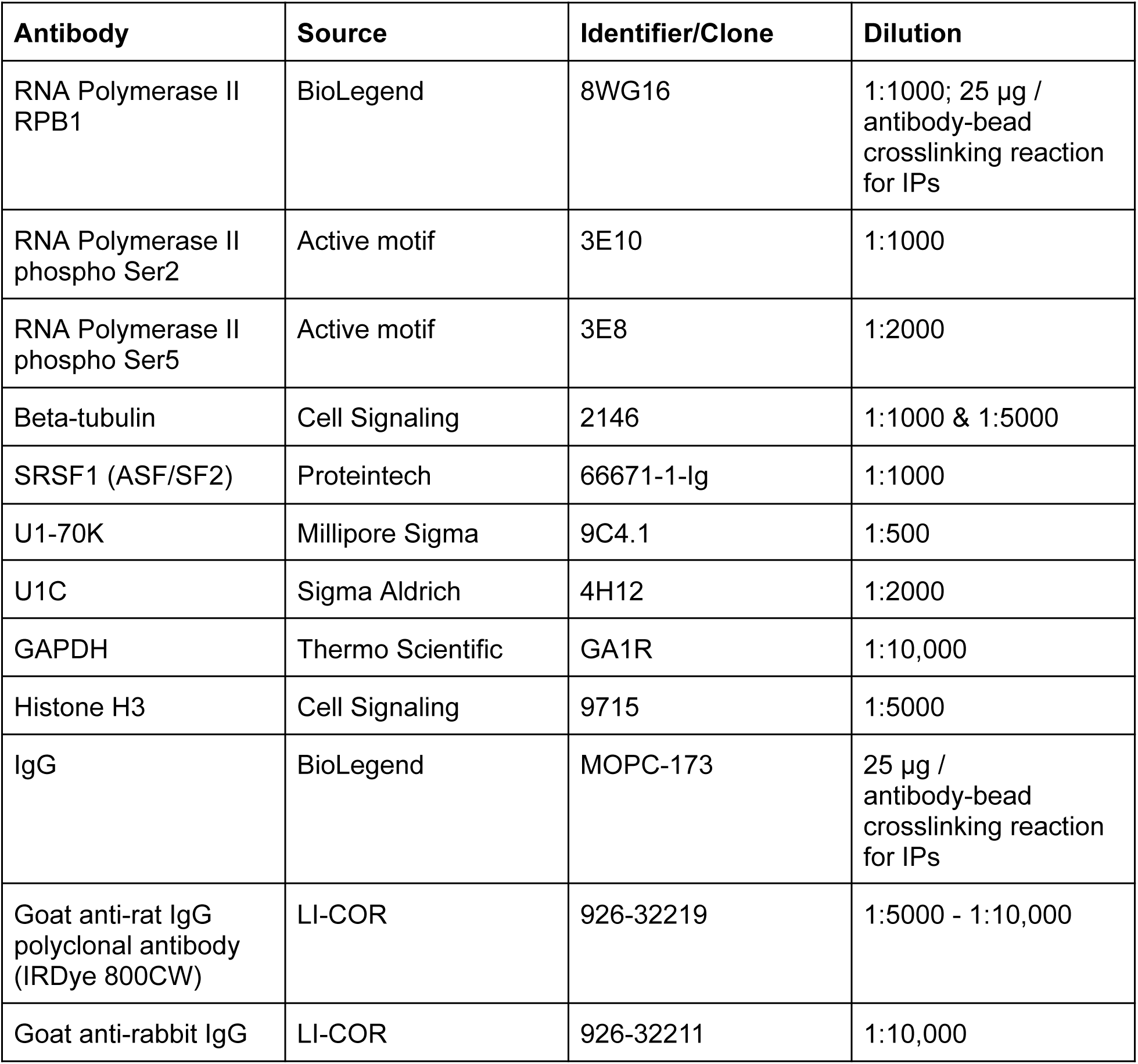
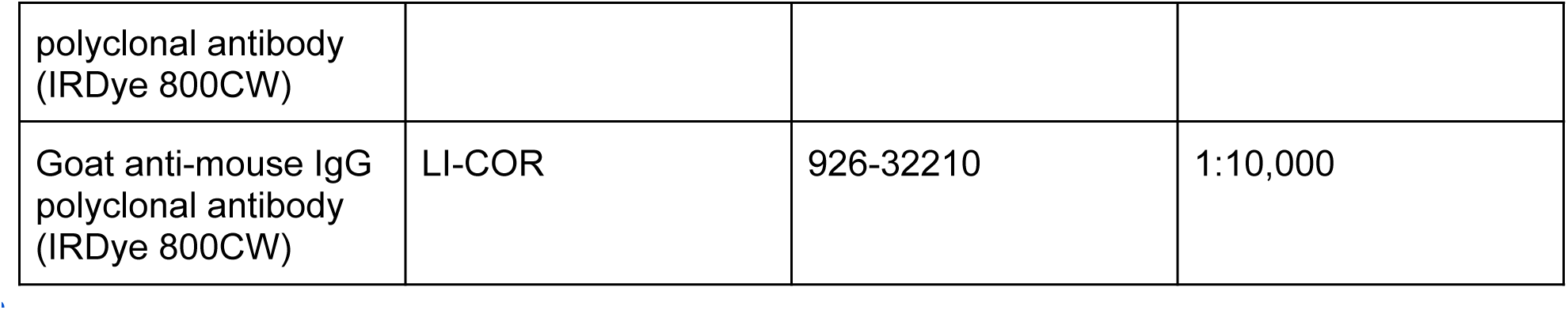

#### Chromatin RNA 3′-end sequencing

To collect chromatin-associated RNA for 3′-end sequencing, target (SRSF1 or U1-70K) or non-silencing control shRNA cell lines were seeded by splitting confluent plates 1:10 (day 0). On days 1-3, media was replaced and doxycycline was added to a final concentration of 1 µg/ml. On day 3, cells were split approximately 1:4 (∼5 million cells) for collection on day 4.

On day 4, one plate of cells (approximately ≤10 million cells) were collected for each cell fractionation reaction. Cell fractionation was performed as described previously^74–76^ and all steps were performed on ice with fresh, pre-chilled buffers. The protocol minorly deviated from previous protocols in the following way: the chromatin pellets used for RNA extraction were resuspended in 200 µL of RIPA buffer (Sigma #R0278) and 750 µL of TRIzol LS (Invitrogen #10296010), rather than QIAzol or RIPA alone. They were resuspended using a 1 mL syringe and 22G needle, and stored at −80°C until RNA extraction.

Chromatin-associated RNA was extracted with TRIzol LS using the protocol recommended by the manufacturer. Samples were resuspended in 180 µL of RNase-free water. RNA yield was measured on a NanoDrop spectrophotometer (ThermoFisher). Any samples with low 260/230 (<1.80) ratios on the NanoDrop were re-precipitated. Samples were then submitted to the Biopolymers Facility at Harvard University for RNA high sensitivity TapeStation analysis.

For DNase treatment, each sample received 20 µL of 10X DNase I buffer and 2 µL of amplification grade DNase I (Invitrogen #18068015). Samples were incubated in the DNase mixture at room temperature for 30 minutes. To re-isolate RNA, phase-lock tubes (QuantaBio #10847-802) were prepared by centrifuging tubes at 14,000 g for one minute at room temperature. RNA samples were then added to each prepared phase-lock tube. One volume of chloroform-isoamyl alcohol (24:1) was subsequently added to each tube, and tubes were mixed by manual shaking for >15 seconds. Samples were centrifuged at 16,000 g for 5 minutes at room temperature, and the upper aqueous phase was transferred to a new microcentrifuge tube. To recover RNA in a starting volume of 150 µL, 1/10 volume of 5M NaCl and 2.5X volumes of 100% ethanol were added. RNA was incubated overnight at −20°C. The following day, RNA was spun down at 4°C for 30 minutes at 20,000 g, and subsequently washed twice with fresh, room temperature 75% ethanol. Following ethanol washes, supernatant was removed, and RNA pellets air dried for a few minutes. RNA pellets were then resuspended in nuclease-free water, heated at 65°C for 2 minutes, and resuspended with a pipette. Final RNA concentrations were measured on Qubit, and samples were submitted to the Biopolymers Facility at Harvard University for RNA high sensitivity TapeStation analysis. Processed samples were stored at −80°C until library preparation.

Chromatin-associated RNA 3′-end sequencing libraries were prepared using the QuantSeq 3′ mRNA-Seq Library Prep Kit REV V2 from Lexogen (#225.24). Input for each sample was ∼500 ng of RNA. Library integrity was assessed with a dsDNA high sensitivity Bioanalyzer assay, TapeStation, and qPCR at the Bauer Core Facility at Harvard University, where libraries were subsequently sequenced with the CSP primer from the Lexogen kit, on the Illumina NovaSeq X Plus platform. Paired-end 100 base pair libraries (n = 3 biological replicates per experiment) were sequenced to a depth of ∼70-182 million reads.

#### 3′-RACE and whole cell RNA-sequencing

Cells were treated with siRNAs against SRSF1 or U1-70K and a non-targeting control for 3 days, as described above. Upon cell collection, plates were rinsed once with room temperature 1X PBS, before being treated with trypsin, quenched with ice cold media (DMEM + 10% FBS + 1% P/S), collected, and spun down at 300 g for 4 minutes at 4°C. Cells were again resuspended in 10 mL ice cold 1X PBS before again being spun down at 300 g for 4 minutes at 4°C. Cells were resuspended in 300 µL RIPA buffer (Sigma #R0278) and 250 µL of sample was used for RNA collection. RNA samples received 750 µL of TRIzol LS. Samples sat for 2-5 minutes at room temperature before being vortexed until homogenous and stored at −80°C until RNA extraction. Whole cell RNA was either extracted with TRIzol LS using the protocol recommended by the manufacturer and DNase-treated as described above for chromatin-associated RNA using amplification grade DNase I (Invitrogen #18068015), or was extracted with TRIzol LS using the miRNeasy Mini Kit (Qiagen #217004) and DNase-treated using the RNase-Free DNase Set (Qiagen #79254).

To validate 3′-end sequencing data, we performed 3′ RACE (Rapid Amplification of cDNA Ends), according to the manufacturer’s protocol (Invitrogen, 18373019). We performed cDNA synthesis using the oligo-dT containing Adapter Primer. The first PCR used a forward primer (outer primer) designed to bind upstream of a known PAS in a gene of interest (either an unaffected PAS or altered termination site identified using 3′-end sequencing), and the reverse primer was the Abridged Universal Amplification Primer (AUAP) from the kit. We performed a second, nested PCR with 2 µL of the first PCR reaction, with a forward primer (nested primer) designed to bind downstream of the first forward primer, along with the same AUAP as the reverse primer. Taq DNA polymerase (NEB M0273) was used for all PCR reactions.

3′ RACE was performed for genes *ANAPC7, IFNAR1,* and *RIOK3*. For *ANAPC7*, 28 cycles were used for each PCR reaction at 52°C annealing temperature. For *IFNAR1*, 22 cycles were used for each PCR reaction at 51°C or 52°C annealing temperature. For *RIOK3*, 22 cycles were used for PCR 1 at 51°C annealing temperature, and 25 for PCR 2 at 49.5°C annealing temperature. Primer sequences are listed below.

3′ RACE gels were quantified using the Image Lab software’s volume tools, using the adjusted volume (intensity) for each amplicon. The proximal PAS/distal PAS intensity was calculated separately for control and KD samples, and relative intensity was compared as a ratio of KD to control for each replicate. A paired t-test was used for determining statistical significance with * indicating p < 0.05.

In addition to 3′ RACE, DNase-treated whole cell RNA was sent for KAPA mRNA HyperPrep library preparation and sequencing at the Bauer Core Facility at Harvard University. 100 base pair paired-end sequencing was performed on an Illumina NovaSeq SP flow cell (n = 3 biological replicates for SRSF1 and control KD samples, n = 2 biological replicates for U1-70K and control KD replicates) to a depth of ∼80-120 million reads per sample.

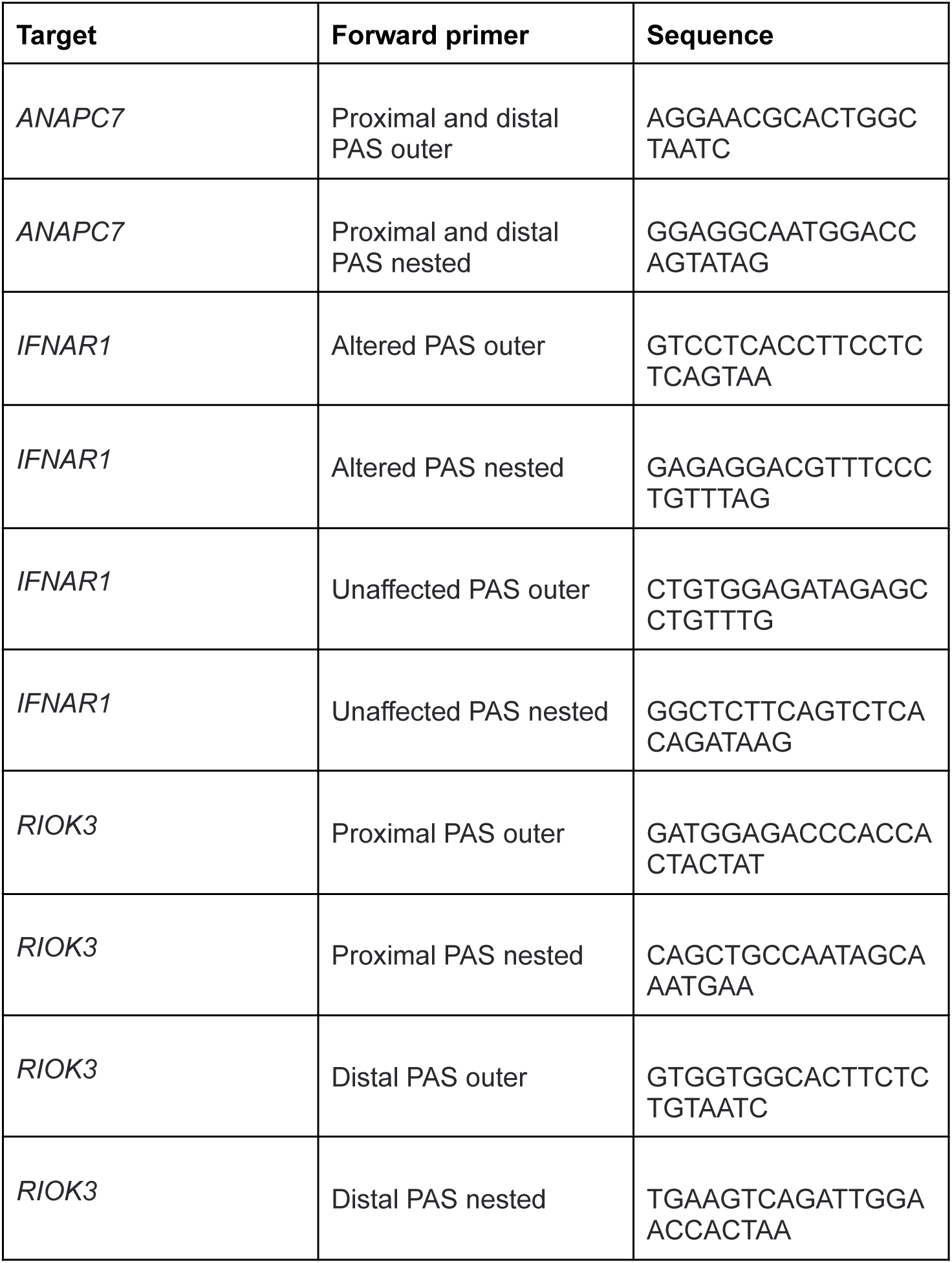

#### Immunoprecipitation-mass spectrometry

For immunoprecipitations (IPs) performed following shRNA-mediated target or non-silencing knockdowns, confluent cells were seeded to ∼1 million cells per plate on the first day of doxycycline treatment (0.5 µg/ml doxycycline used for U1-70K knockdowns, 1 µg/ml doxycycline used for SRSF1 knockdowns). For the next two days, cells received fresh media and doxycycline each day. Three days following the initial doxycycline treatment, cell lysates were collected for IP.

To perform dimethyl pimelimidate (DMP) crosslinking of antibodies to beads, 62.5 µL of Protein A Sepharose beads (GE healthcare #17-5280-04) were washed three times with 1X PBS. For each crosslinking reaction, 25 µL of Pol II antibody (Biolegend 8WG16) and 25 µL of 1X PBS were added to beads; for control bead sets, 50 µL of IgG antibody (Biolegend MOPC-173) was added to beads. Beads and antibodies were coupled by rotating at room temperature for one hour. Following one hour, supernatant was removed, and coupled beads and antibodies were washed twice with 1 ml of 0.2 M sodium borate (pH 9) (diluted from VWR #AAJ63637-AK or made from Sigma #S9640). Coupled beads and antibodies were stored in 1 ml of 0.2 M sodium borate (pH 9) solution while a fresh solution of 20 mM DMP (Sigma D8388 or Thermo Scientific 21667) in 0.2 M sodium borate (pH 9) was made. Coupled beads and antibodies were spun down and supernatant was removed. 1.5 ml of the 20 mM DMP solution was added to each crosslinking reaction, and crosslinking occurred through rotation at room temperature for 30 minutes. Following crosslinking, the supernatant was removed, and crosslinking was stopped by washing twice with 1 ml of 0.2 M ethanolamine (pH 8) (Sigma E6133). Samples were rotated at room temperature for 2 hours in 1 ml of 0.2 M ethanolamine (pH 8). Samples were then washed twice with 1X PBS. Finally, beads were resuspended in 970 µL of 1X PBS and NaN_3_ was added to a final concentration of 0.3%. Crosslinked beads and antibodies were stored at 4°C until use. Supernatant was removed for each crosslinking step with a 25G 1.5 inch long needle and each spin was performed at 500 g for 2 minutes at 4°C. All steps were performed on ice unless otherwise noted.

Nuclear lysates were prepared for IP as in Folco *et al.* 2012^77^, with the following modifications: the concentration step prior to dialysis was omitted, and for dialysis, 50 µL of extract was added to each Slide-A-Lyzer MINI device (10K MWCO, Thermo #69572) and was dialyzed in a slowly stirred dialysis buffer for 2 hours at 4°C.

Pol II IPs were performed as previously described, with modifications^54,78^. A reaction mixture was made up that was comprised of 30% HEK293T nuclear extract, 30% splicing dilution buffer (diluted from stock of 20 mM HEPES (pH 7.9) and 100 mM KCl), 500 µM ATP, 3.2 mM MgCl_2_, and 20 mM phosphocreatine di(tris) salt, for a total volume of either 500 µL or 333 µL, depending on the experiment. The reaction mixture was incubated for 20 minutes at 30°C. IP buffer (1X PBS, 0.1% Triton-X 100, 0.6 mM PMSF, 1X protease inhibitor (Roche #04693132001)) was added such that it constituted approximately one-third of the final volume, and the mixture was spun down at 13,000 rpm for 5 minutes. The supernatant was recovered to use as nuclear IP lysate. IP input samples were taken from the recovered nuclear lysate (approximately 6.67% of the lysate volume). Prepared crosslinked beads and antibodies were spun down for 2 minutes at 500 g at 4°C and their storage solution was removed. Nuclear lysate was rotated with crosslinked beads and antibodies overnight on a nutator mixer at 4°C. If conducting AMO experiments using the U1 AMO (GGTATCTCCCCTGCCAGGTAAGTAT, Gene Tools) or scrambled control AMO (CCTCTTACCTCAGTTACAATTTATA, Gene Tools), then 12 µM of the AMO was added to the respective lysate, bead, and antibody mixture prior to overnight rotation.

Following overnight rotation, beads were washed five times with 1.5 mL of wash buffer (1X PBS, 0.1% Triton-X 100, 0.2 mM PMSF). Beads in the wash buffer were placed in a new tube using a cut tip before the final spindown in order to remove background proteins. After the last wash, the wash buffer was removed completely. Proteins were eluted by adding a volume of protein gel buffer (125 mM Tris (pH 6.8), 5% SDS, 20% glycerol, 0.005% bromophenol blue) of approximately 12-16% of the volume of the nuclear lysate input to the IP. In the protein gel buffer, beads were gently vortexed, spun down, gently vortexed again, and then incubated for 20 minutes at room temperature. Beads were spun down for 2 minutes at 500 g at room temperature, and the final elute was recovered. Following elution, DTT was added to samples at a final concentration of 40 mM.

To prepare samples for mass spectrometry, proteins were precipitated from the protein gel buffer (Calbiochem #539180) and submitted to the Taplin Mass Spectrometry Facility at Harvard Medical School for processing.

#### Transient transcriptome sequencing (TT-seq)

TT-seq was performed as previously reported^79,80^. Briefly, cells were treated with siRNAs against SRSF1 and a non-targeting control for 3 days, as described above. All steps were performed with minimal light exposure, to protect 4sU from light. Prior to cell collection on day 4, 4sU (Sigma T4509 or Cayman Chemical Company #16373) was added to fresh media to a final concentration of 500 µM. For 4sU labeling, media was aspirated from plates and replaced with an equal volume of pre-warmed 4sU-containing media. Plates were placed in the 37°C incubator for a 12 minute 4sU-labeling period. Cells were then washed once with room temperature 1X PBS before 2 mL of TRIzol LS (Invitrogen #10296010) was directly added to plates. Plates were stored at room temperature for 3 minutes before cells were scraped into Eppendorf tubes. The samples were pipetted and vortexed to homogenize cells completely before being placed on ice prior to RNA extraction or being stored at −80°C.

Additional reference plates were also collected, which received identical siRNA treatments as the experimental plates and were collected for whole cell protein extraction for western blots. Reference plate cells were rinsed once with room temperature 1X PBS, before being treated with trypsin, quenched with ice cold media (DMEM + 10% FBS + 1% P/S), collected, and spun down at 300 g for 4 minutes at 4°C. Cells were again resuspended in 10 mL ice cold 1X PBS before again being spun down at 300 g for 4 minutes at 4°C. Cells were resuspended in 300 µL RIPA buffer (Sigma #R0278) and 50 µL of sample was used for protein collection. To protein samples, 1 µL of benzonase (Sigma #E1014) and 1 µL of 50X protease inhibitor solution (Roche #04693132001) were added. Samples were then vortexed briefly and incubated on ice for 30 minutes. Samples were then spun down at maximum speed for 10 minutes at 4°C before collecting the supernatant for western blotting.

4sU-labeled RNA was extracted with TRIzol LS using the protocol recommended by the manufacturer. Samples were resuspended in 50 µL of RNase-free water. Samples within the same condition were combined (2 tubes) for a total of 100 µL of RNA per condition. RNA yield was measured on a NanoDrop spectrophotometer. Any samples with low 260/230 (<1.80) ratios on the NanoDrop were reprecipitated.

TT-seq library construction was performed by the Nascent Transcriptomics Core at Harvard Medical School, Boston, MA. As part of library construction, 4sU-labeled *Drosophila* RNA was added as a spike-in to samples. TT-seq samples were sequenced on Illumina NovaSeq 6000 and NextSeq 500 platforms with 42-111 base pair paired-end reads to a depth ∼70-167 million reads/sample. Three independent biological replicates were sequenced; one replicate consisted of two pooled biological samples.

#### Precision run-on sequencing (PRO-seq)

After 3 days of siRNA KD of SRSF1 or U1-70K and non-targeting controls, ∼6-8M cells were collected for cell permeabilization, as previously described^81^. Trypan blue was used to determine permeabilization efficiency, with samples with >95% permeabilization used for library preparation. Samples were flash frozen in liquid nitrogen, and stored at −80°C. Whole cell protein was extracted from reference plates for western blots.

PRO-seq library construction was performed by the Nascent Transcriptomics Core at Harvard Medical School, Boston, MA, as previously reported^81^. As part of library construction, *Drosophila* RNA was added as a spike-in to samples. PRO-seq samples were sequenced on Illumina NextSeq 2000 and NovaSeq X+ platforms with 50 base pair paired-end reads following basemasking to a depth ∼24-50 million reads/sample. Three independent biological replicates were sequenced per experiment.

### QUANTIFICATION AND STATISTICAL ANALYSIS

#### Chromatin RNA 3′-end sequencing data analysis

FASTQ data was trimmed using bbduk with settings k=13 ktrim=r useshortkmers=t mink=5 qtrim=t trimq=10 minlength=20, and aligned with STAR v2.7.9a using the following settings: –outSAMmode Full –outSAMattributes Standard –outFilterMismatchNmax 16 –outFilterMultimapScoreRange 1 –outfilterMultimapNmax 10 –outSAMunmapped None –outReadsUnmapped Fastx –outSAMmapqUnique 255. Sam files were sorted, indexed, and converted to bam files using Picard Tools, and bam files were further filtered for uniquely mapping reads (samtools view -q255 -hb).

Samples were analyzed using a custom pipeline. Part one of the chromatin RNA 3′-end sequencing data analysis pipeline consisted of making a reference file of all the PASs recovered from the 3′-end sequencing data using custom scripts. To do this, bam files were first converted to bed files using BedTools bam2bed^82^ and then PASs were identified and clustered. Briefly, from the input bed files, the first base of read 1 was isolated to define 3′-end site coordinates. Next, to filter out reads potentially stemming from genomic polyA stretches, the sequence of the 3′-end site and 10 downstream base pairs was analyzed and sequences containing >7 As were removed. Reads were filtered for those aligning to non-snRNA or snoRNA containing protein coding genes using Bedtools intersect (wo=True, s=True)^82^. 3′-end sites within 25bp of each other were clustered together as a single site (Bedtools clusters, s=True, d=25), and read sums within each cluster were used as PAS counts downstream. The outer boundaries of the 3′-end sites within each cluster were used as the start and end sites for each PAS cluster and were expanded with an additional 5 base pairs at each boundary. To make a GTF-formatted reference file of high confidence PAS coordinates, 3′-end site clusters were filtered for those with ≥20 counts. The remaining clusters were combined across samples for each genomic strand respectively using Bedtools multiinter to annotate intervals of PAS coordinates shared across replicates. Intervals populated by ≥2 samples were kept, and subsequently, directly adjacent (book-ended) intervals were joined using Bedtools merge in a strand-aware manner. Bedtools map was used to add back hg38 gene name and attribute information to each corresponding PAS interval in a strand-aware manner; every interval mapped to a gene had to map to that gene uniquely in order for that gene and its respective PAS intervals to be retained for downstream analysis. The resulting annotations of PAS intervals were converted into a GTF file for downstream use.

Part two of the nascent RNA 3′-end sequencing data analysis pipeline consisted of conducting differential expression analysis on the 3′-end sequencing data, and using the 3′-end site GTF created above to do so. First, the 3′ positions of uniquely mapped bam file reads were extracted using a custom script. Counts for 3′-end sites were calculated using featureCounts, using the custom PAS GTF as the reference file, and useMetaFeatures=False. To only retain high confidence 3′-end sites, we filtered sites for those with at least 50 counts in at least one condition, and we furthermore filtered to only consider genes with more than one PAS. To evaluate the differential usage of PASs relative to gene expression, we used the package DEXseq^83^ and plotted results in R.

#### Browser track analyses

For 3′-end sequencing samples, bigwig files of the first base of read 1 (i.e. the 3′-end position of each read) were extracted from uniquely mapped bam files using deeptools^84^ bamCoverage with the following settings: -of bigwig --normalizeUsing CPM --Offset 1 --samFlagInclude 64 --binSize 1 for --filterRNAstrand forward and --filterRNAstrand reverse. Browser track analyses of these bigwig files were performed and plotted using pyGenomeTracks^85,86^. 3′-end sequencing data tracks were uniformly thickened for visualization in Illustrator.

Browser track analyses of TT-seq data were performed on spike-in normalized bigwig files made from uniquely aligned bam files and plotted using pyGenomeTracks^85,86^.

#### Genomic feature mapping analyses

Genomic features (exons, introns, 5′ UTRs, and 3′ UTRs) were derived and extracted from *Homo sapiens* Ensembl annotations version 108 using makeTxDbFromEnsembl(), exonsBy(), intronsByTranscript(), fiveUTRsByTranscript(), and threeUTRsByTranscript() from the GenomicFeatures package^87^. Genomic features were mapped to PASs using the annotate_regions() function from the annotatr package^88^. To avoid assigning the same PAS to multiple genomic features, PASs mapped to more than one genomic feature were assigned to a single feature using the following prioritization scheme: 5′ UTR, 3′ UTR, exon, intron. Feature prioritization was not used for upset plots. Pie charts were plotted using ggplot2^89^ and upset plots were made using the ComplexUpset package^90^.

To map the positions of PASs within 3′ UTRs, Ensembl gene and 3′ UTR annotations were retrieved for *Homo sapiens* using biomaRt^91^. Annotations for overlapping or adjacent 3′ UTRs within the same gene and on the same strand were reduced using IRanges, 3′ UTRs were ordered and numbered in a strand-aware manner from 5′ to 3′, and PASs were mapped to 3′ UTRs using findOverlaps with maxgap = −1^92,93^.

#### TCGA data retrieval

TCGA data used in this study was curated by The Cancer Genome Atlas (TCGA) Research Network^51^ and obtained from dbGaP under accession phs000178. Breast tumor sample data was retrieved that had previously been categorized as having high or low SRSF1 expression levels compared to diploid samples^52^. Expression level z-scores for samples were used as pre-calculated by cBioPortal^94,95^.

#### HITindex pipeline analysis

The hybrid-internal-terminal (HIT) index pipeline was run as reported previously^46^ for SRSF1 and control knockdown whole cell RNA-seq data to characterize and measure relative alternative last exon (ALE) usage (<u>github.com/thepailab/HITindex</u>). The identified ALEs were then run through the HITindex statistical analysis pipeline (<u>github.com/fiszbein-lab/HIT_Index_Stat_Analysis-</u>) to identify significantly differentially expressed last exons across conditions (|percent spliced in (PSI)| > 0.1 and false discovery rate (FDR) < 0.05).

For TCGA breast tumor sample analysis, we compared the two populations of tumors expressing different levels of SRSF1, rather than analyzing paired measurements between two conditions. Therefore, for each tumor, we calculated an overall ALE bias score (distal ALE - proximal ALE usage) for each gene. Each per-tumor bias score was calculated as the median of ALE bias scores across genes.

#### RNA binding site analyses

We obtained Protein-RNA Interaction with dual-deaminase Editing and Sequencing (PIE-seq) RNA binding site data from Ruan *et al.* 2023^47^, which characterized RNA binding protein (RBP) binding patterns for SRSF1, PUM2, and NOVA1 in HEK293FT cells.

We used deeptools^84^ to make bigwig files of PAS data and to map the PAS signal around SRSF1 and other RBP PIE-seq binding sites. For datasets of all PAS sites, we started with sorted, uniquely mapped bam files of chromatin-associated 3′-end sequencing data and used bamCoverage -of bigwig --normalizeUsing CPM --Offset 1 --samFlagInclude 64 --binSize 1 for --filterRNAstrand forward and --filterRNAstrand reverse. For datasets subsetted by the effect of SRSF1 knockdown on PAS usage, we began with the 3′ positions of uniquely mapped bam file reads of chromatin-associated 3′-end sequencing data and used bamCoverage -of bigwig--normalizeUsing CPM --Offset 1 --binSize 1 for --filterRNAstrand forward and --filterRNAstrand reverse. Next, to compute 3′-end sequencing coverage matrices around identified PIE-seq binding site coordinates within the 3′ UTR, deeptools computeMatrix was used with the bigwigs generated and the following settings: computeMatrix reference-point -b 2000 -a 2000 --averageTypeBins “sum” --missingDataAsZero --binSize 1.

Metagene plots of averaged PAS signal around binding sites were generated using a custom script. Matrices generated from full chromatin-associated 3′-end sequencing datasets were used for PAS signal, which had proper counts per million (CPM) normalization in bamCoverage. To get subsetted data, these full datasets were filtered for corresponding PAS signal within matrices subsetted by the effect of SRSF1 knockdown on PAS usage. Binding sites were filtered for those with surrounding PAS signal, and then further filtered for those with the top 75% highest surrounding PAS signal within 2 kb. Replicates were averaged together. To account for global changes in gene expression, we further analyzed the relative PAS signal around binding sites, dividing each matrix bin by the total sum of signal around each binding site. Finally, to look at the metagene signal, for each position around the binding site, PAS signal was averaged across binding sites and the standard error was calculated.

#### Immunoprecipitation-mass spectrometry data analysis

For IP-MS analysis (n = 4), log_2_ transformed sum intensity values were either median value or bait normalized. For analyses of Pol II IPs with U1 or scrambled AMO treatments, NA values were randomly imputed for each condition from a distribution with its mean and standard deviation based on the bottom 10% of normalized values. For analyses of Pol II vs. IgG IPs, NA values were randomly imputed across all conditions from the bottom 10% of normalized values^96,97^. For each protein, a Welch’s t-test was used to determine if significant differences existed between conditions. Log_2_ fold change data was plotted in R. GO terms were determined relative to genome-wide annotation for *H. sapiens* (org.Hs.eg.db) in R using clusterProfiler^98^.

#### AlphaFold3 structure predictions

SRSF1 and U1-70K full-length translated sequences were obtained from NCBI Consensus CDS (CCDS) and used with AlphaFold3^53^ to predict SRSF1 binding to U1-70K. As phosphomimetic SRSF1 mutants have been shown to bind U1-70K more readily than wild-type^41^, the 12 serines between residues 205 and 227 were modeled with phosphorylation. Predicted aligned error (PAE) was used to assess the presence of a predicted SRSF1 - U1-70K binding interface. The AlphaFold3 predicted phosphorylated SRSF1/U1-70K structure was mapped to the cryoEM structure of a transcribing Pol II and U1 snRNP^32^ (PDB: 7B0Y) in ChimeraX and residues are shown as indicated in the figure legend.

#### TT-seq data analysis

TT-seq FASTQ files were aligned to a concatenated human GRCh38 and *Drosophila* BDGP6 genome using STAR (v2.5.4a)^99^. STAR settings were as followed: --outSAMmode Full, --outSAMattributes Standard, --outFilterMultimapScoreRange 1, --outFilterMultimapNmax 10, --outSAMunmapped None, --outReadsUnmapped Fastx, and --outSAMmapqUnique 255. Reads were sorted and filtered for uniquely mapping reads.

For differential expression analysis, featureCounts (GTF.featureType=’gene’, useMetaFeatures=TRUE, ignoreDup=TRUE, strandSpecific=2, isPairedEnd=TRUE, requireBothEndsMapped=TRUE, checkFragLength=TRUE, minFragLength=50, maxFragLength=600, autosort=TRUE), was used to assign gene counts to either a modified human or *Drosophila* genome from uniquely aligned reads^100^. The modified human genome used for the featureCounts analysis was filtered for non-overlapping nuclear-encoded protein coding genes that did not contain annotated snRNA or snoRNAs. Sample counts were subsequently used in DESeq2^101^ analysis for differential expression, and were normalized by *Drosophila* spike-in-derived size factors, as estimated by estimateSizeFactors() for *Drosophila* gene counts. Genes with an adjusted p-value < 0.05 and absolute log_2_ fold change greater than log_2_(1.5) between knockdown and control conditions were considered to have significantly changed nascent expression.

For metagene analysis, stranded bigwig files were created from uniquely mapped reads using the deepTools function bamCoverage^102^. Scaled matrices were created using the computeMatrix scale-regions function with the following settings: -b 2500 -m 10,000 -a 2500 –skipZeros --missingDataAsZero and using the same modified human genome as described for the featureCounts analysis above. The resulting matrices for each sample were normalized by their respective *Drosophila* spike-in-derived size factors, as calculated by DESeq2. Spike-in normalized gene counts were summed across all replicates and scaled matrix positions, and genes with total counts in the lowest quartile were filtered out of the analysis. Normalized and filtered gene matrices were then averaged across replicates. Mean signal was derived by averaging across all genes for each scaled position.

#### PRO-seq data analysis

PRO-seq data was processed using an established pipeline^81^. UMIs were extracted using UMI tools, and adapters were removed from reads using cutadapt -f fastq -O 1 –match-read-wildcards -m 26. Reads were aligned with bowtie2 to a concatenated human and *Drosophila* genome in order to map reference and spike-in reads respectively, which were subsequently separated from each other. Samples were deduplicated using UMI tools. Paired end reads were separated from each other in order to isolate out reads indicating the 5′ or 3′ end of the aligned read. Using the custom script bowtie2stdBedGraph.py (-x -a -o b -b 10 -D -l, https://github.com/AdelmanLab/NIH_scripts), PRO-seq samples were converted to bedgraph files.

For analyses of genes with significantly altered PRO-seq signal, the 3′ positions of aligned, deduplicated PRO-seq bam file reads were extracted using a custom script. Read counts for the *Drosophila* spike-in and human protein coding genes were calculated using featureCounts^100^. Significantly altered genes were identified using DESeq2^101^ with *Drosophila* spike-in-based size factor normalization and the results were plotted in R.

#### Pol II elongation index analysis

Pol II elongation index calculations were performed as reported previously^33,79,103^. Using a custom script, bam files of TT-seq data were sorted and indexed using samtools^104^ before making non-normalized bedgraphs with STAR 2.7.9a^99^ with the following command: STAR --runMode inputAlignmentsFromBAM --outWigType bedGraph --outWigStrand Stranded --outWigNorm None. Subsequently, *Drosophila* spike-in reads were removed from bedgraphs, which were further filtered for reads from human chromosomes 1-22, X, Y. PRO-seq bedgraphs were made as detailed above.

Next, for each set of TT-seq or PRO-seq replicate data, each replicate was normalized by its respective size factor. The size factors were generated based on mapping human reads with featureCounts^100^ to human protein coding genes, *Drosophila* reads with featureCounts to the *Drosophila* genome, and deriving *Drosophila* size factors using DESeq2^101^. The dominant transcription start site (TSS) and transcription end site (TES) coordinates for human protein coding genes were determined by GetGeneAnnotation (<u>github.com/AdelmanLab/GetGeneAnnotation_GGA</u>) from PRO-seq and RNA-seq data of control, SRSF1, and U1-70K knockdown samples. Protein coding genes overlapping snRNAs, snoRNAs, or other protein coding genes on the same strand were filtered out from the annotation using a custom script.

Finally, following size factor normalization, replicate samples were merged in a strand-aware manner. TT-seq samples were converted to bigwigs using the UCSC tool bedGraphToBigWig and then merged using the UCSC tool bigWigMerge^105^. PRO-seq samples were merged using bedgraphs2stdBedGraph (github.com/AdelmanLab/NIH_scripts).

Merged bedgraphs were next used to make binned position-based matrices with the make_heatmap script (github.com/AdelmanLab/NIH_scripts/). We looked at genomic positions downstream of the TSS using 500 basepair bins for TT-seq or PRO-seq data.

To process TT-seq and PRO-seq matrices of protein coding genes for Pol II elongation index calculations, a custom script was used. Briefly, this script removed bins that were upstream, overlapping, or 500 basepairs downstream of the TSS, as well as bins at and downstream of the TES. TT-seq bins were multiplied by their average fragment length (300 basepairs), and a pseudocount of 1 was added to all datasets to avoid dividing by zero. The TT-seq and PRO-seq bins with the lowest 5% counts were filtered out as low confidence bins. The elongation index was calculated as a ratio of TT-seq reads per bin to the PRO-seq reads per bin.

For metagene plots of the elongation index, the mean elongation index across genes for each positional bin was calculated with a 95% confidence interval. For boxplots, the log_2_ elongation index values for each bin were plotted according to knockdown condition.

#### Readthrough index analysis

Protein coding genes considered for the readthrough analysis were filtered using a custom script that started with gene annotations from the dominant TSS to TES. These dominant sites were as determined by the GetGeneAnnotation (GGA) (<u>github.com/AdelmanLab/GetGeneAnnotation_GGA</u>) analysis, based on PRO-seq and RNA-seq data of control, SRSF1, and U1-70K knockdown samples. Protein coding genes overlapping snRNAs, snoRNAs, and other protein coding genes on the same strand were filtered out from the annotation; additionally, protein coding genes with downstream regions that overlapped snRNAs, snoRNAs, lncRNAs, or other protein coding genes within 20 kilobases on the same strand were also filtered out. Only genes over 1 kilobase were kept in the analysis. To assign last exon coordinates to dominant TESs, exon coordinates from the Homo sapiens Ensembl release version 108 were retrieved using makeTxDbFromEnsembl from the GenomicFeatures package^87^ and mapped to dominant TESs using the annotatr package^106^. If a dominant TES mapped as the 3′ end to more than one exon within a gene, the longest exon was chosen; last exons had to be at least 50 basepairs to be retained for analysis.

To calculate the readthrough index, the last exon coordinates for the dominant TES were first identified and paired with the 20 kilobase downstream region coordinates for that gene. TT-seq featureCounts^100^ data for last exons and downstream regions was normalized by DESeq2-derived size factors^101^, based on GGA annotations, as described above, and a pseudocount of 1 was added to each normalized dataset. Last exons and downstream regions were filtered for those surpassing a minimum count threshold, and genes were filtered for those where the DESeq2 log_2_ fold change in expression was < 0.15. Finally, counts were length normalized prior to calculating the average counts across replicates within each last exon or downstream region per gene. The average readthrough index was calculated as the ratio of downstream reads to last exon reads. Additionally, readthrough indices were calculated for individual replicates. Genes considered for further analysis had a readthrough index <= 0.8 in the control condition. Reproducible readthrough events were those for which all three individual replicates for a gene had a readthrough index ≥1.2. Readthrough indices were plotted in R.

## Supplementary Tables

**Table S1:** IP-mass spectrometry-derived fold change values in Pol II co-IP with factors after treatment with a U1-targeting or scrambled AMO.

## Resource availability Lead contact

Further information and requests for resources and reagents should be directed to and will be fulfilled by the lead contact, Stirling Churchman.

## Materials availability

Unique and stable reagents generated in this study are available upon request.

## Acknowledgements

We thank J. Warner, I. Patop, F. Winston, and L. Hansen for helpful comments and critical reading of the manuscript. We thank R. Reed and B. Chi for assistance in troubleshooting the Pol II IP from nuclear lysate experiments. We thank the Harvard Bauer Sequencing Core for assistance with library preparation and RNA sequencing experiments. We thank S. Goldman and the Harvard Nascent Transcriptomics Core for advice and assistance with building and sequencing TT-seq and PRO-seq libraries and discussing data analysis. We thank R. Tomaino from the Taplin Mass Spectrometry Facility at Harvard University and E. McShane for assistance with IP-mass spectrometry experiments. We thank the Harvard Bioinformatics Core for advice on statistical analyses. We thank the Biopolymers Facility at Harvard University for running samples for Sanger sequencing and TapeStation. Additionally, we thank the Adelman lab for helpful discussions on elongation index analyses. We thank S. Sedor and A. Elchert for helpful advice regarding AlphaFold3 analyses. We thank G.Y. Kim for discussions on the HITindex pipeline analysis. We thank C. Chang for experimental support. H.E.M. was supported by a National Science Foundation Graduate Research Fellowship DGE 1745303. This work was supported by NIH grants (F32GM15606301 to A.M.R, R01GM136794 and R61CA309729 to L.S.C., and R35GM147254 to A.F.).

## Author contributions

H.E.M. and L.S.C. conceived of and designed the experiments. H.E.M. performed the experiments. For U1-70K knockdown chromatin-associated RNA 3′-end sequencing samples, H.E.M. and A.M.R. performed cell fractionations and western blots, and A.M.R. prepared the libraries. A.M.R. collected PRO-seq samples and ran western blots. H.E.M., A.M.R., and M.K. performed 3’ RACE assays. H.E.M. performed data analysis. C.L.C. analyzed whole cell RNA-seq data and cancer samples from the TCGA database with the HITindex pipeline. H.E.M., A.F., and L.S.C. interpreted the results. H.E.M., A.M.R., A.F., and L.S.C. wrote the manuscript with input from all authors.

## Declaration of generative AI and AI-assisted technologies in the manuscript preparation process

During the preparation of this work the authors used ChatGPT, Claude, and Grammarly in order to assist in editing the manuscript. After using these tools, the authors reviewed and edited the content as needed and take full responsibility for the content of the published article.

---

**Supplementary Figure 1.**
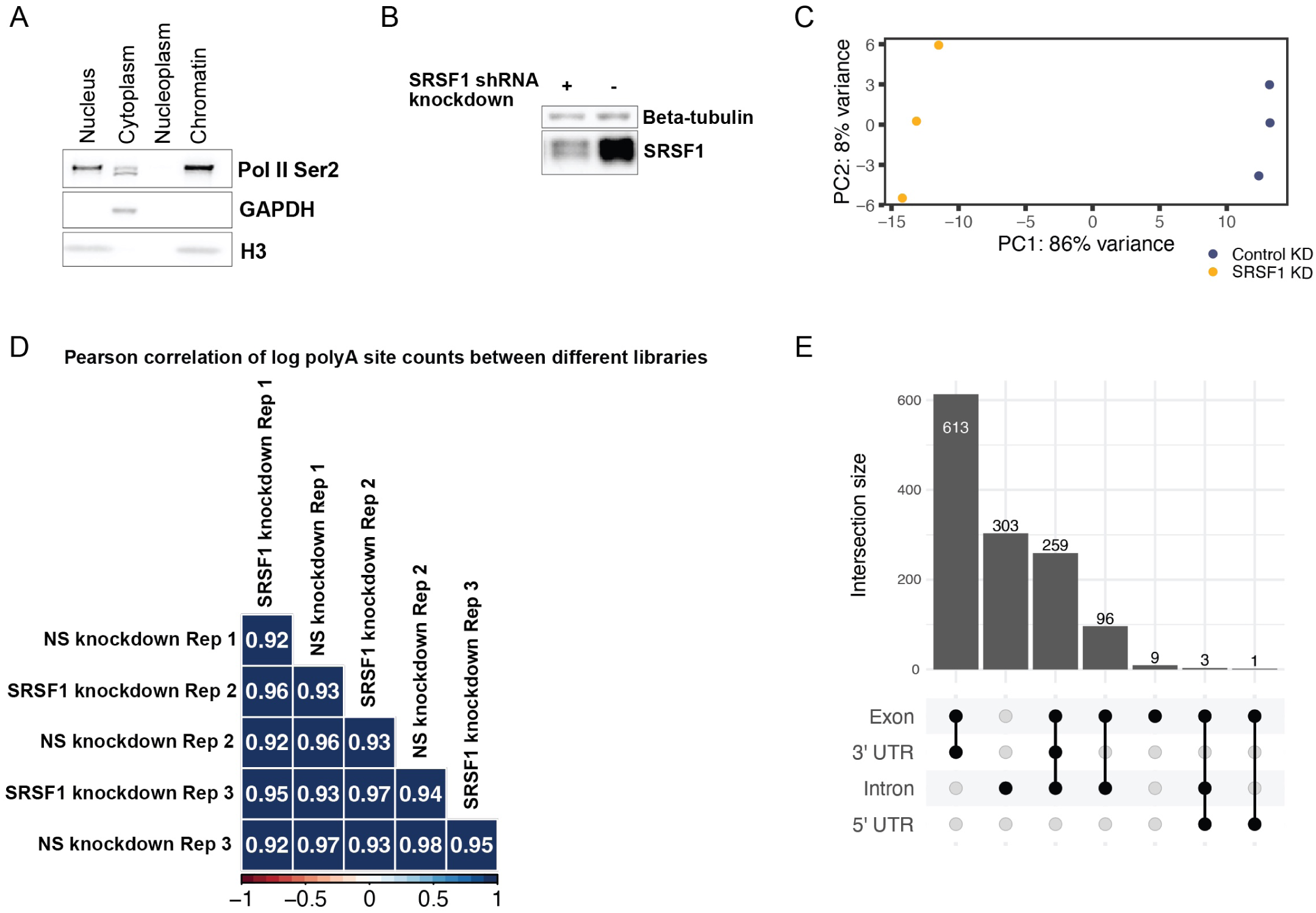
SRSF1 regulates PAS selection. A) Western blot of subcellular fractionation of HEK293T cells. GAPDH signal marks the cytoplasmic fraction, and enriched Pol II Ser2 and H3 signal mark the nuclear and chromatin fractions. One representative replicate shown. B) Western blot showing SRSF1 KD (+) with shRNAs; in the control condition (-), expression of a non-silencing (NS) shRNA was induced. C) PCA plot for SRSF1 KD 3′-end sequencing samples. PCA performed based on PAS counts. D) Pearson correlation of log_2_(PAS counts) biological replicates from SRSF1 and control KD 3′-end sequencing samples. Sites with 0 counts across all samples were filtered out and a pseudocount of 1 was added to each site. E) Upset plot of the feature annotations in Fig. 1G with no prioritization for feature assignment.

**Supplementary Figure 2.**
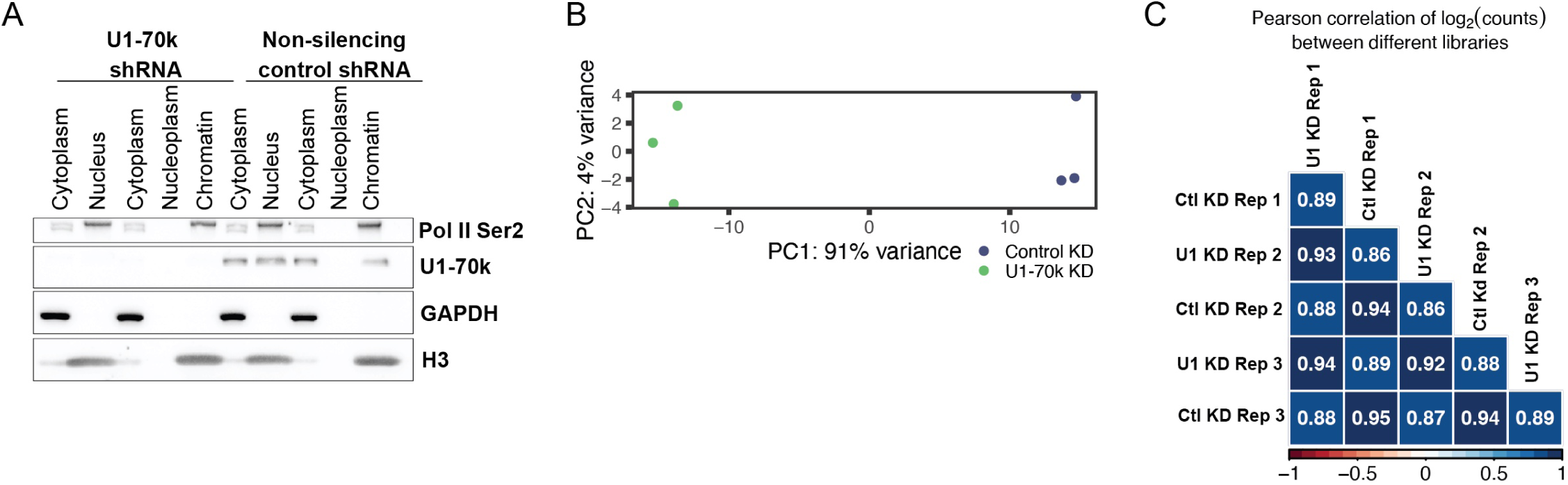
SRSF1 and U1 snRNP regulate a shared set of PASs. A) Western blot of subcellular fractionation of HEK293T cells for U1-70K shRNA KD experiments. GAPDH signal marks the cytoplasmic fraction, and enriched Pol II Ser2 and H3 signals mark the nuclear and chromatin fractions. One representative replicate shown. B) PCA plot for U1-70K KD, chromatin-associated RNA 3′-end sequencing samples. PCA performed based on PAS counts. C) Correlation of log_2_ PAS counts across U1-70K and control (Ctl) KD, chromatin-associated RNA 3′-end sequencing samples. Sites with 0 counts across all samples were filtered out and a pseudocount of 1 was added to each site.

**Supplementary figure 3.**
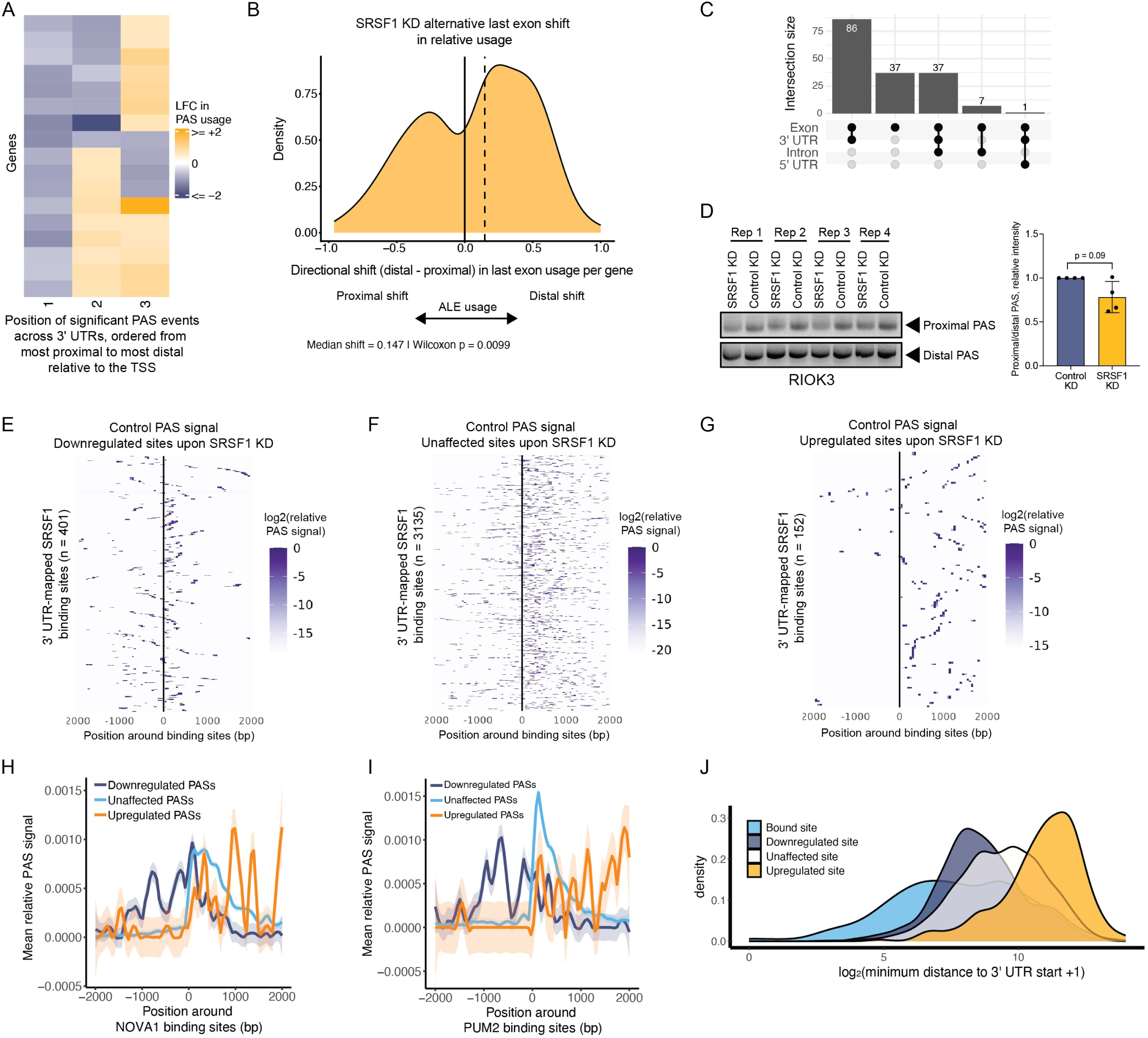
SRSF1 binds towards the start of 3′ UTRs, near SRSF1-sensitive PASs. A) Heatmap of the log_2_ fold change (LFC) in PAS usage across one or more 3′ UTRs with 3 significantly altered proximal/distal PASs (n = 17). Sites ordered from most proximal to most distal. Changes in PAS usage were clustered by Euclidean distance and Ward.D linkage using ComplexHeatmap. B) HITindex pipeline analysis of the overall directional shift (distal - proximal) in ALE usage for each gene (n = 207) between SRSF1 KD and control whole cell RNA samples. Analysis based on events with significant changes in ALE usage (|delta PSI| > 0.1, FDR < 0.05). The dotted line indicates the median shift; the solid line indicates the median value if the overall shift were 0. C) Upset plot of the annotations generated in Fig. 3D without feature prioritization. D) 3′ RACE assay performed on whole cell RNA upon SRSF1 or control KD (left) and quantification (right) for *RIOK3* (n = 4). E-G) Heatmaps of the combined control sample (n = 3) relative PAS signal +/-2 kb around 3′ UTR-mapped SRSF1 binding sites. Heatmaps subsetted by the effects on the PASs following SRSF1 KD. The relative usage of the 75% highest subsetted PAS signal was plotted. The data shown here was averaged over all rows within each heatmap for the metagene plot in Fig. 3G. H-I) Metagene plots, plotted as in Fig. 3G, around PIE-seq RNA binding sites for NOVA1 (H) and PUM2 (I). J) 3′ UTRs containing both PIE-seq SRSF1 binding sites and PASs or only PASs were assessed for the log_2_ distance (in basepairs (bp), +1 bp pseudocount) distributions between the start of each 3′ UTR and the nearest binding site or PAS, with PASs subsetted by the effect of SRSF1 KD.

**Supplementary figure 4.**
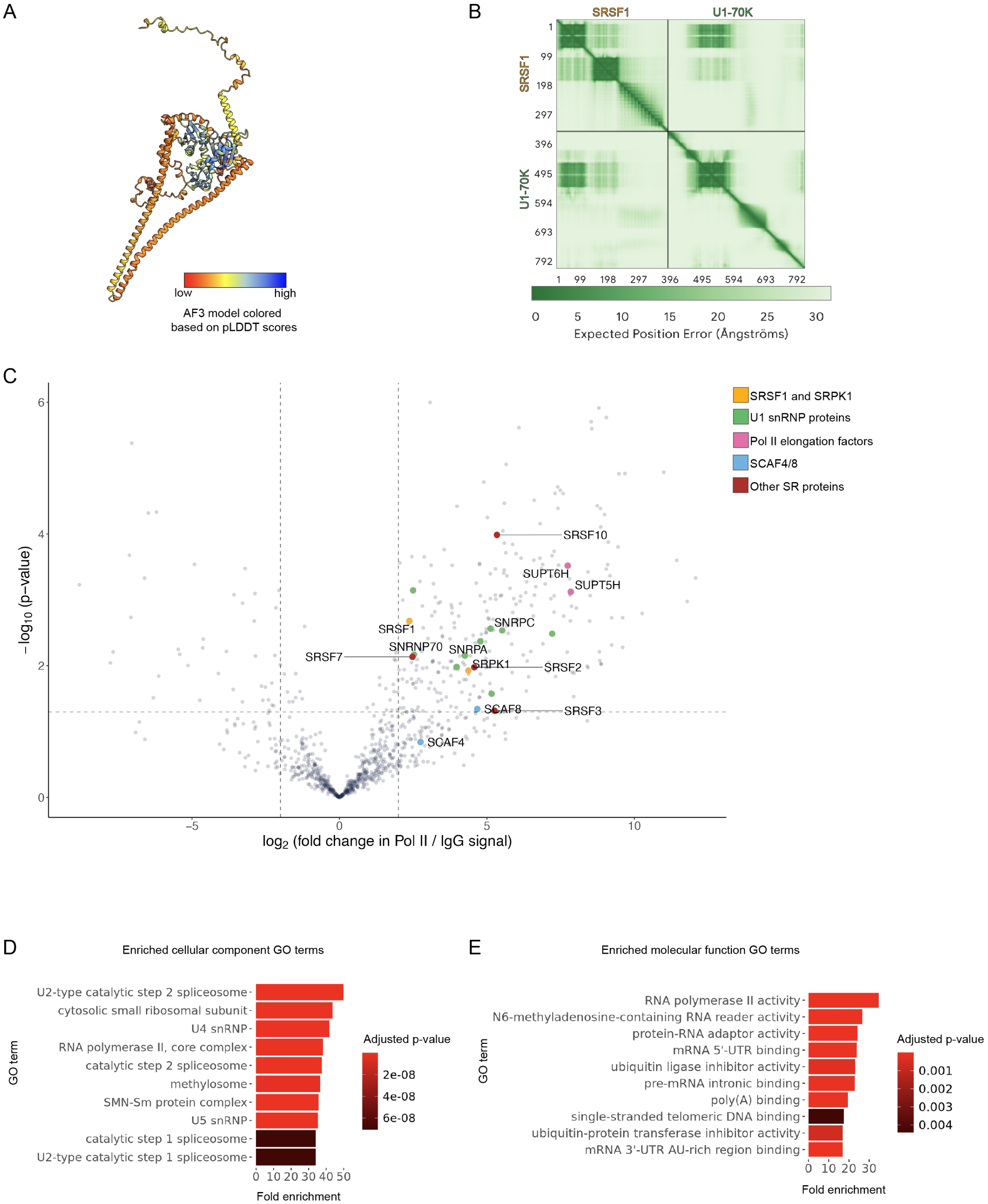
Interactions between SRSF1 and Pol II are dependent on U1 snRNP. A) AF3 structure of full-length phosphorylated SRSF1 and U1-70K. Colored by predicted local distance difference test (pLDDT) scores. B) Predicted aligned error (PAE) plot for the phosphorylated SRSF1-U1-70K AF3 structure. C) Volcano plot of the enriched proteins in Pol II vs. IgG IP-MS samples (n = 4). All samples treated with the scrambled (control) AMO. Significance threshold set at |log_2_ fold change for Pol II IP enrichment| ≥2 and -log_10_ p-value ≥1.3. D-E) Cellular component (D) and molecular function (E) gene ontology (GO) terms for proteins significantly enriched for Pol II IP over IgG IP as shown in C.

**Supplementary Figure 5.**
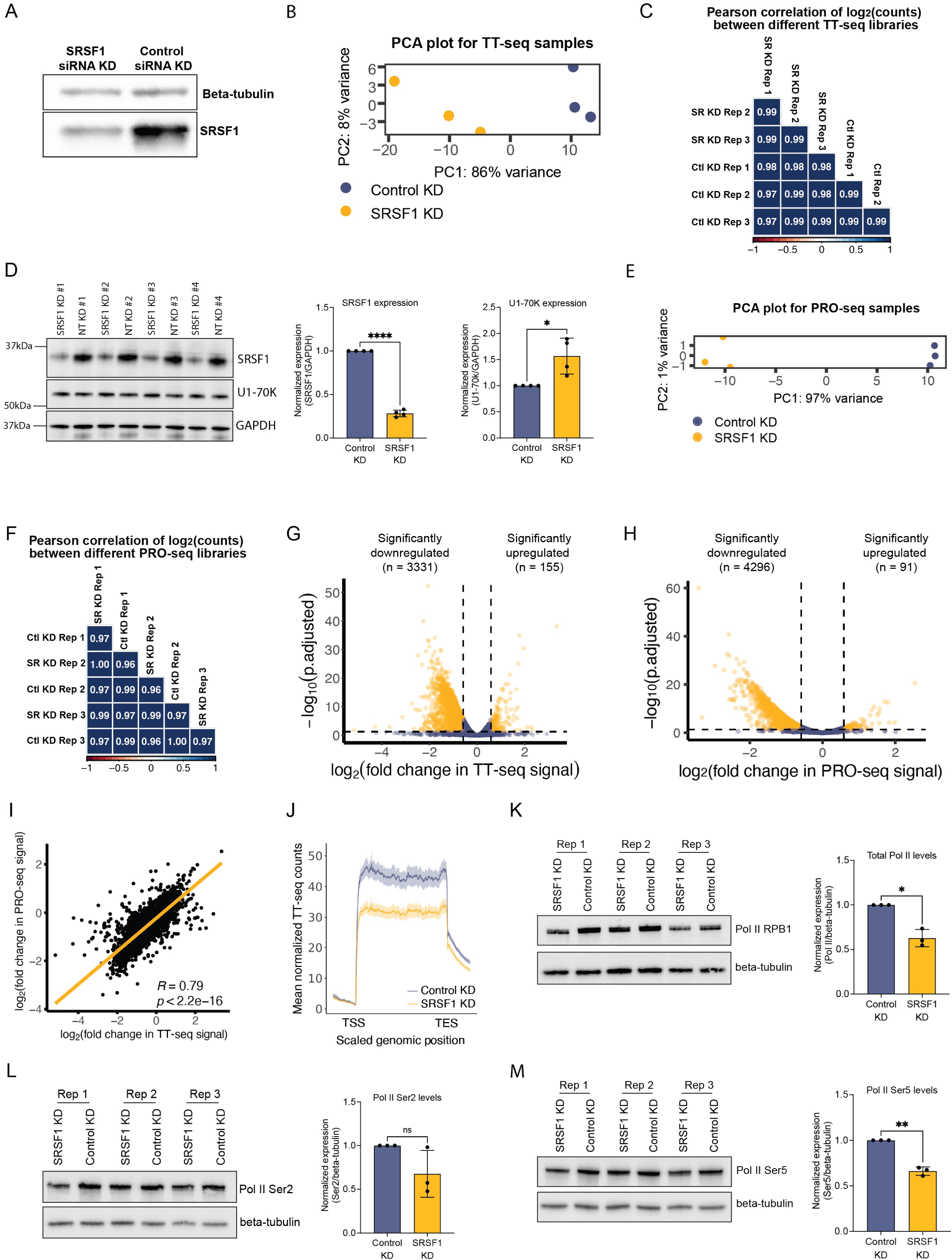
SRSF1 slows Pol II elongation and limits transcription readthrough. A) Western blot of total protein samples for the representative TT-seq experiment shown. Samples were treated either with a SRSF1-targeting siRNA or non-targeting control (NT) siRNA for 72 hours. Beta-tubulin served as a loading control. B) PCA analysis of TT-seq data for SRSF1 KD samples. C) Pearson correlation analysis of biological replicates from SRSF1 KD and control TT-seq samples using spike-in normalized counts. D) Western blot (left) of total protein reference samples for PRO-seq experiments. Samples were treated either with a SRSF1-targeting siRNA or non-targeting control (NT) siRNA for 72 hours. GAPDH served as a loading control for SRSF1 and U1-70K KD quantification (right). E) PCA analysis of PRO-seq samples aligned to protein-coding genes for SRSF1 KD samples. F) Pearson correlation analysis of biological replicates from SRSF1 KD and control PRO-seq samples using spike-in normalized counts. G) Volcano plot of nascent gene expression changes (from TT-seq data) upon SRSF1 KD compared to control samples. Data normalized using *Drosophila* spike-in reads; significant events thresholded with p.adjusted < 0.05, |log_2_ fold change| > log_2_(1.5). Upregulated genes: 155, downregulated genes: 3331, unaffected genes: 7358. Each dot is one protein-coding gene. Analysis performed with DESeq2. H) Volcano plot of changes in PRO-seq signal upon SRSF1 KD compared to control samples. Data normalized using *Drosophila* spike-in reads; significant events thresholded with p.adjusted < 0.05, |log_2_(fold change)| > log_2_(1.5). Upregulated genes: 91, downregulated genes: 4296, unaffected genes: 6476. Each dot represents one protein-coding gene. Analysis performed with DESeq2. I) Pearson correlation calculation between the log_2_ fold change in TT-seq signal and the log_2_ fold change in PRO-seq signal. Log_2_ fold change values calculated between SRSF1 and control KD samples. J) SRSF1 and control KD TT-seq signal scaled metagene plot for the top 75% highest expressed protein-coding genes, with a +/- 95% confidence interval. n = 6549 genes in the SRSF1 KD condition, n = 6516 genes in the control KD condition. K-M) Western blots and quantifications of total protein samples treated either with a SRSF1-targeting siRNA or non-targeting control siRNA for 72 hours. Samples probed for total Pol II levels using a CTD-targeting RPB1 antibody (K), Pol II Ser2 (L), and Pol II Ser5 (M). Beta-tubulin served as a loading control.

